# Centrosome impairment causes DNA replication stress through MLK3/MK2 signaling and R-loop formation

**DOI:** 10.1101/2020.01.09.898684

**Authors:** Zainab Tayeh, Kim Stegmann, Antonia Kleeberg, Mascha Friedrich, Josephine Ann Mun Yee Choo, Bernd Wollnik, Matthias Dobbelstein

## Abstract

Centrosomes function as organizing centers of microtubules and support accurate mitosis in many animal cells. However, it remains to be explored whether and how centrosomes also facilitate the progression through different phases of the cell cycle. Here we show that impairing the composition of centrosomes, by depletion of centrosomal components or by inhibition of polo-like kinase 4 (PLK4), reduces the progression of DNA replication forks. This occurs even when the cell cycle is arrested before damaging the centrosomes, thus excluding mitotic failure as the source of replication stress. Mechanistically, the kinase MLK3 associates with centrosomes. When centrosomes are disintegrated, MLK3 activates the kinases p38 and MK2/MAPKAPK2. Transcription-dependent RNA:DNA hybrids (R-loops) are then causing DNA replication stress. Fibroblasts from patients with microcephalic primordial dwarfism (Seckel syndrome) harbouring defective centrosomes showed replication stress and diminished proliferation, which were each alleviated by inhibition of MK2. Thus, centrosomes not only facilitate mitosis, but their integrity is also supportive in DNA replication.

**Highlights:**

- Centrosome defects cause replication stress independent of mitosis.
- MLK3, p38 and MK2 (alias MAPKAPK2) are signalling between centrosome defects and DNA replication stress through R-loop formation.
- Patient-derived cells with defective centrosomes display replication stress, whereas inhibition of MK2 restores their DNA replication fork progression and proliferation.

**Graphical abstract:** 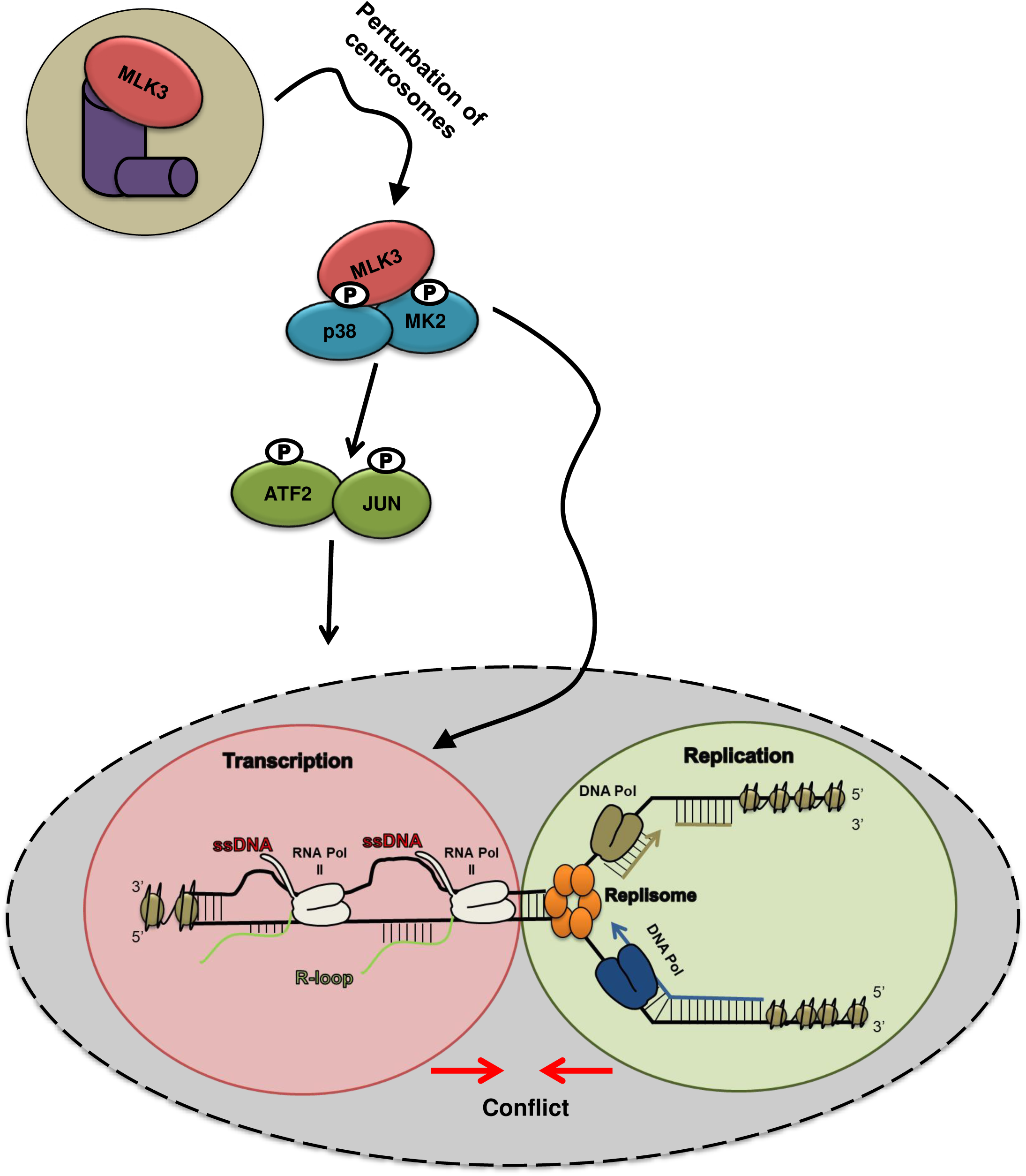

## BACKGROUND

The centrosome is a large complex of proteins, built around the centrioles and mostly known for its function as a microtubule organizing center. This not only serves to shape a major component of the cytoskeleton, but also to form the cilia. Another important role for centrosomes appears from their association with the poles of the mitotic spindle (Nigg and Raff, 2009). Current evidence suggests that centrosomes support the accurate distribution of sister chromatids towards the two daughter cell nuclei during mitosis (Nam et al., 2015). However, the precise role of centrosomes in this context is still under debate. Fungal and plant cells do not require centrosomes for their division, and even in animals, not all cells employ them in mitosis. Thus, the precise function(s) of centrosomes are less obvious than initially assumed, justifying a search for additional activities performed by them.

The term “Seckel syndrome” is commonly used to describe autosomal recessive conditions mainly characterized by primary microcephaly, intellectual disability, short stature, and a typical facial gestalt (Griffith et al., 2008). Two seemingly distinct groups of gene defects have been associated with this condition.

On the one hand, hypomorphic recessive alleles of *ATR* (Ataxia telangiectasia and Rad3 related) or the functionally associated gene *ATRIP* lead to Seckel syndrome (O’Driscoll et al., 2003). In parallel, malfunction, inhibition or depletion of ATR strongly impairs the replication of DNA (Flynn and Zou, 2011). The kinase ATR and its substrate CHK1 are each required for smooth progression of DNA replication forks, and compromising their function results in a condition termed replication stress (Dobbelstein and Sorensen, 2015). This stress condition is characterized by decreases in replication fork progression, ultimately resulting in DNA damage and a corresponding, ATR-driven signaling cascade.

On the other hand, morphological traits that overlap with Seckel syndrome are also found in association with mutations in genes encoding centrosomal components (Kalay et al., 2011; Zhou et al., 2014). Microcephalic primordial dwarfism was also described in patients harbouring mutations in *PLK4* (Dinçer et al., 2017), a known master regulator of centrosome biogenesis (Kim et al., 2013; Dinçer et al., 2017). At least at first glance, it seems puzzling why mutations affecting either the DNA replication machinery, or otherwise the centrosome, result in clinically similar phenotypes.

A reminiscent coincidence was found in mouse models. Genetic disruption of ATR led to Seckel-like symptoms in mice (Murga et al., 2009). Again, the same was observed in mice carrying deletions in the centrosomal CEP63 gene product (Marjanovic et al., 2015). Being aware that a number of reasons might impair the proliferation of neuronal cells in these models, this still led us to test whether the integrity of the centrosome, like ATR, might sustain proper DNA replication.

In our previous work, we have identified MAPKAPK2 alias MK2 as a key kinase that mediates replication stress (Kopper et al., 2013) at least in part by phosphorylating and counteracting translesion DNA polymerases. MK2 is part of a signalling pathway that transmits stress signals from the upstream kinases summarized as p38 (MAPK14, MAPK11, MAPK12, and MAPK13) to its targets, and it forms a stable complex with p38 (Gaestel et al., 2006).

Here, we show that the disruption of centrosome integrity induces replication stress. This is true even when the cells were not allowed to go through mitosis while impairing the centrosomes. Centrosome damage induces a specific signaling cascade reaching from centrosome-associated MLK3, through p38 and MK2, to the transcription factors ATF2 and JUN. This results in the formation of RNA:DNA hybrids (R-loops), causing replication stress. These observations strongly suggest that centrosomes not only have a role in mitosis, but that their integrity is also required for the unperturbed progression of DNA replication forks.

## Materials and Methods

Materials are listed in detail in Supplementary Table 1.

### Cell culture

H1299 (non-small cell lung carcinoma, p53 -/-), SW48 (colon carcinoma) and retinal pigment epithelial (RPE) cells, as well as patient-derived skin fibroblasts were maintained in Dulbecco’s modified Eagle’s medium (DMEM). One set of fibroblasts was derived from a microcephaly patient carrying a homozygous mutation of CEP152, i.e. the splice site mutation c.261+1G>C. This mutation and its consequences were previously described (Kalay et al., 2011). The mutation gives rise to several splice variants, many of which result in a frame shift. However, one of them leads to an in frame deletion of only two amino acid residues. Moreover, when overexpressed, this mutant localizes to centrosomes. These observations suggest that the mutant is hypomorphic and still capable of expressing some functional CEP152, but not at the level of its wildtype counterpart. Cell culture media were supplemented with 10% fetal calf serum (FCS) and antibiotics (penicillin, streptomycin, and ciprofloxacin). Cells were maintained at 37°C in a humidified atmosphere with 5% CO2. All cell lines were routinely tested for mycoplasma contamination and scored negatively.

### Treatments and transfections

Stock solutions of small compounds were prepared in DMSO and then diluted in pre-warmed medium. For each siRNA-knockdown, cells were reverse transfected with 10nM siRNA (Ambion/Thermo Fisher), using Lipofectamine 3000 (Invitrogen). Culture media were changed after 24 hrs, followed by further incubation for 24 hrs. Plasmid transfections were carried out using 2µg of plasmid DNA with Lipofectamine 2000. Media were changed after 6 hrs followed by incubation for 48 hrs.

### Fiber assays

DNA fiber assays to analyze replication fork progression and processivity were essentially carried out as described (L A Tibbles, 1996). Following drug treatment or transfection with pooled siRNAs and/or plasmid DNA, cells were incubated with 5-chloro-2′-deoxyuridine (CldU), 25μM for 20 min, followed by 5-iodo-2′-deoxyuridine (IdU; both from Sigma-Aldrich), 25μM for 60 min. DNA fibers were spread and lysed on glass slides using a spreading buffer (200mM Tris pH 7.4, 50mM EDTA, 0.5% SDS). After acid treatment (2.5M HCl), CldU- and IdU-labeled tracks were detected by 1 hr incubation with rat anti-BrdU antibody (1:400, AbD Serotec; 1:1000, Abcam; detects BrdU and CldU) and mouse anti-BrdU antibody (1:400; detects BrdU and IdU; Becton Dickinson). Slides were fixed in 4% paraformaldehyde in phosphate buffered saline (PBS) and incubated for 2 hrs with Alexa Fluor 555-conjugated goat anti-rat antibody and Alexa Fluor 488-conjugated goat anti-mouse antibody (both 1:200; Invitrogen). Fiber images were acquired by fluorescence microscopy and analyzed manually using Image J, assuming that a track length of 1 μm corresponds to 2.59 kb. Two-tailed Mann-Whitney t tests and Student’s t-tests were calculated by Graph Pad Prism 6 and 7.

### R-loop detection (immunofluorescence)

Cells were grown on coverslips overnight prior to transfection or treatment, washed once with PBS, and fixed with 100% cold methanol for 10 minutes at −20 °C, three brief washes with PBS containing 0.1% Tween 20 (PBST) and blocking with 3% bovine serum albumin (BSA) in PBST for 1 hr. The primary antibody S9.6 (Absolute antibody) was diluted in the blocking buffer (1:100) and incubated overnight at 4 °C, followed by incubation with 488 Alexa Fluor-coupled donkey anti mouse IgG (H+L) antibody (Invitrogen) 1:250 for 2 hrs at room temperature. Coverslips were washed with PBST and were briefly incubated with 1:2000 4′,6-diamidino-2-phenylindole (DAPI), followed by mounting (DAKO). Fluorescence signals were detected by a microscope (Zeiss Axio Scope.A1) equipped with filters for 488nm, a EC Plan-Neofluar 100x oil objective and an Axiocam 503 color camera. Per condition, approximately 15-20 images were taken with the AxioVision software and analyzed using ImageJ.

### R-loop detection (dot blot analysis)

H1299 cells were fixed with 1.1% PFA in Buffer A (0.1M NaCl, 1 mM EDTA, 0.5 mM EGTA, 50 mM HEPES) for 30 min, followed by quenching with glycin. The cells were then harvested in Buffer B (0.25% Triton X-100, 10 mM EDTA, 0.5 mM EGTA, 20 mM HEPES), resuspended in Buffer C (0.15 M NaCl, 1 mM EDTA, 0.5 mM EGTA, 50 mM HEPES) and then in IB Buffer (0.5% Triton X-100, 0.375% SDS, 0.75 M NaCl, 5 mM EDTA, 2.5 mM EGTA, 100 mM HEPES). Upon sonication (Bioruptor, Diagenode, 10 cycles, 30 sec each) and centrifugation, the supernatant was treated with proteinase K at 50°C for 1 hr, phenol-extracted and ethanol-precipitated. 500ng of the DNA in 2 µL H2O was spotted onto pre-wet nitrocellulose membrane and cross-linked with UV-C. The membrane was blocked with 5% BSA in PBS-T (0.25% Tween-20) and incubated with the monoclonal antibody S9.6 (1:300 in 5% BSA/PBS-T) for 16 hrs at 4°C, followed by secondary antibodies. For normalization and in parallel, the DNA was denatured with 2.5 M HCl for 10 min, washed with PBS-T and incubated with antibody to single stranded DNA (ssDNA; 1:1000) followed by secondary antibody. The peroxidase-coupled secondary antibodies were detected by luminescence.

### Immunofluorescence analysis of centrosomes

20000 cells were seeded on 8-well chamber slides (Nunc/Thermo, cat #177445) overnight prior to treatment. Cells were treated with drugs or siRNA for 48 hrs. The slide was washed once with phosphate buffered saline (PBS), fixed and stained as above. Antibodies were used as follows: 1:200 PCNT (Abcam), 1:50 CEP152 (Sigma), 1:100 MLK3 (Abcam), 1:500 of 488 Alexa fluor Donkey anti mouse, 1:500 594 Alexa fluor goat anti rabbit, 1:2000 Hoechst 33342.

### Flow cytometry

Cells stained for DNA content with propidium iodide were analyzed using a Guava® Express Pro EasyCyte flow cytometer and Cyto Soft 5.3 software (Merck). Measurements were taken by counting 10000 events at 300-700 cells/mL. To determine the number of gated cells in each cell cycle phase (G1, S, G2/M), histogram markers were adjusted to the profile of asynchronous cells.

### Cell proliferation assay (Celigo)

Cells were seeded at a density 5*10^3^ cells/well in 24-well plates and treated with DMSO or Centrinone. Their proliferation capacity was measured using the Celigo Cytometer (Nexcelom, software version 2.0) by automated light microscopy. Cell confluence was measured every 24 hrs for up to 7 days. Experiments were carried out in 2 biological replicates and with 6 technical replicates for each time point.

### Detection of cells in S phase by EdU incorporation

H1299 cells were incubated with 10µM EdU for 2 hrs. Cells were fixed with 3.7% PFA in PBS for 30 minutes and permeabilizated with 0.5% Triton X-100 for 30 minutes. Staining and detecting incorporated EdU was done using the Click-iT EdU Alexa Fluor imaging kit (Invitrogen/Molecular Probes).

### Chromosome spread analysis and chromosome counting

H1299 cells were treated with 2 µM Dimethylenastron (DME, Sigma) for 4.5 hrs, to trap cells in mitosis. The cells were then trypsinized and incubated in 750 µL hypotonic solution (40% medium in water) for 10-15 min. 250 µL of Carnoy’s fixative solution (methanol and glacial acetic acid, 3:1) was added to the pellet. The cells were then resuspended twice in Carnoy’s fixative and stored at −20 °C for 16 hrs. After resuspension in acetic acid, 10 µl of cell suspension was dropped on cold glass slides from ∼ 2 m height. Slides were placed on a 42°C heat block for ∼ 10-15 min for drying and then stained with 8% Giemsa solution for 15 min. The analysis of chromosomes was carried out by transmission microscopy.

### Chromatin fractionation

Cells were synchronized using the CDK4 inhibitor Palbociclib and released into a thymidine block. 48 hrs after treating the cells with 500 nM Centrinone, the cells were trypsinized and collected in 15ml tubes, washed twice with PBS, and centrifuged at 100g for 5 minutes. The pellets were resuspended in 1 ml Buffer A (10mM HEPES, PH 7.9, 10mM KCL, 1.5mM MgCl2 0.34M Sucrose 10% Glycerol, 1mM dithiothreitol and protease inhibitors cocktails (Complete Roche) and transferred into 1.5 ml tubes. Triton X-100 was added to each tube to a final concentration of 0.1%, followed by incubation on a rotating wheel for 15 minutes at 4°C. The tubes were centrifuged at 1300g for 5 minutes at 4°C. The supernatant (cytoplasmic fraction) was transferred to new tubes. The pellets were washed once with Buffer A and then further lysed with 250 μl modified RIPA buffer (1mM EDTA, 150mM NaCl, 0.1% Na-DOC, 1% NP-40, 50Mm Tris pH7.5 and Complete Roche protein inhibitors). 50U of benzonase (nuclease; Novagen) was added to the samples and incubated for 5-10 minutes at room temperature. Samples were mixed by pipetting during the incubation time until they lost viscosity. Samples were diluted 1:3 with modified RIPA buffer. The supernatant was cleared by centrifuging the samples for 7 min at 16000g, 4°C, and the clear chromatin fraction was transferred to a new tube. After boiling the samples in Laemmli buffer at 95°C for 5 min, equal amounts of protein samples were separated by SDS-PAGE, transferred onto nitrocellulose, and stained with the following antibodies: JUN (Santa Cruz), ATF2 (Cell signaling), MCM7 (Cell signaling), GAPDH (Abcam).

### Immunoblot analysis

Cells were harvested in protein lysis buffer (20 mM Tris-HCl pH 7.5, 150 mM NaCl, 1 mM Na2EDTA, 1 mM EGTA, 1 mM beta-glycerophosphate, 2 M urea, proteinase inhibitors pepstatin, leupeptin hemisulfate, aprotinin). The samples were briefly sonicated to disrupt DNA-protein complexes. Total protein concentration was measured using a Pierce BCA protein assay kit (Thermo Scientific Fisher). After boiling the samples in Laemmli buffer at 95°C for 5 min, equal amounts of protein samples were separated by SDS-PAGE, transferred onto nitrocellulose, and visualized with antibodies to the following proteins: MLK3 (D-11) (SantaCruz), PLK4 (Proteintech), p-p38 (E-1) (SantaCruz), p-MK2 (Cell signaling), pHsp27 S82 (Cell signaling), GAPDH (Abcam), RNAseH1 (Abcam), HSC70 (SantaCruz), H3K27me3 (Abcam), H3K27ac (Abcam), Histone H3 (Abcam), P-JUN (S63) (Cell signaling), JUN (Abcam), TBP (SantaCruz), ATF2 (SantaCruz), Phospho-ATF2 (T71) (Cell signaling), RNA Polymerase II (N-20) (SantaCruz), MCM7 (D10A11) XP (Cell signaling), MAPKAPK-2 (Cell signaling), P53 (Cell signaling), phospho-H2A.X (Ser139) (Cell signaling).

## Results

### Depletion of centrosomal components or interfering with PLK4 activity each reduces DNA replication fork progression

To clarify how centrosome integrity affects DNA replication, we inhibited Polo-like Kinase 4 (PLK4) by the small compound Centrinone (Wong et al., 2015), to deplete centrosomes from H1299 cells (lung adenocarcinoma). To assess DNA replication in this context, we performed fiber assays and measured the progression of single replication forks using several cell types. PLK4 inhibition not only led to a strong reduction in the number of detectable centrosomes (Figure 1 A, B and Supplementary Figure 1 A, B), as described (Wong et al., 2015), but also impaired the progression of DNA replication forks (Figure1 C, D and Supplementary Figure 1 C). Similarly impaired fork progression was observed in SW48 cells (colon carcinoma; Figure 1 E and Supplementary Figure 1 E) and in HCT116 cells (colon carcinoma) with or without a deletion of the tumor suppressor TP53 (Figure 1 F, Supplementary Figure 1 F-I). Reduced fork progression upon PLK4 inhibition was also seen in retinal pigment epithelial cells (RPE) that had been immortalized (but not transformed) by hTert (Figure 1 G, Supplementary Figure 1 J). Next, we induced the disintegration of centrosomes by depleting single components, to see how this would affect DNA replication. H1299 cells displayed a reduction in the numbers of detectable centrosomes upon depletion of several centrosomal components by siRNA (Figure 1 H, Supplementary Figure 1 K-N). This was true when depleting constituents of different centrosomal regions, i.e. CEP192 (localized at the inner layer of the pericentriolar matrix, PCM), CEP152 (outer PCM), CCP110 (subdistal appendages), and SASS6 (centrioles) (Supplementary Figure 1 K-N). Remarkably, each of these depletions significantly decreased fork progression (Figure 1 I, Supplementary Figure 1 O). The PLK4 inhibitor CFI-400945, structurally unrelated to Centrinone, impaired DNA replication as well (Supplementary Figure 1 D).

**Figure 1.**
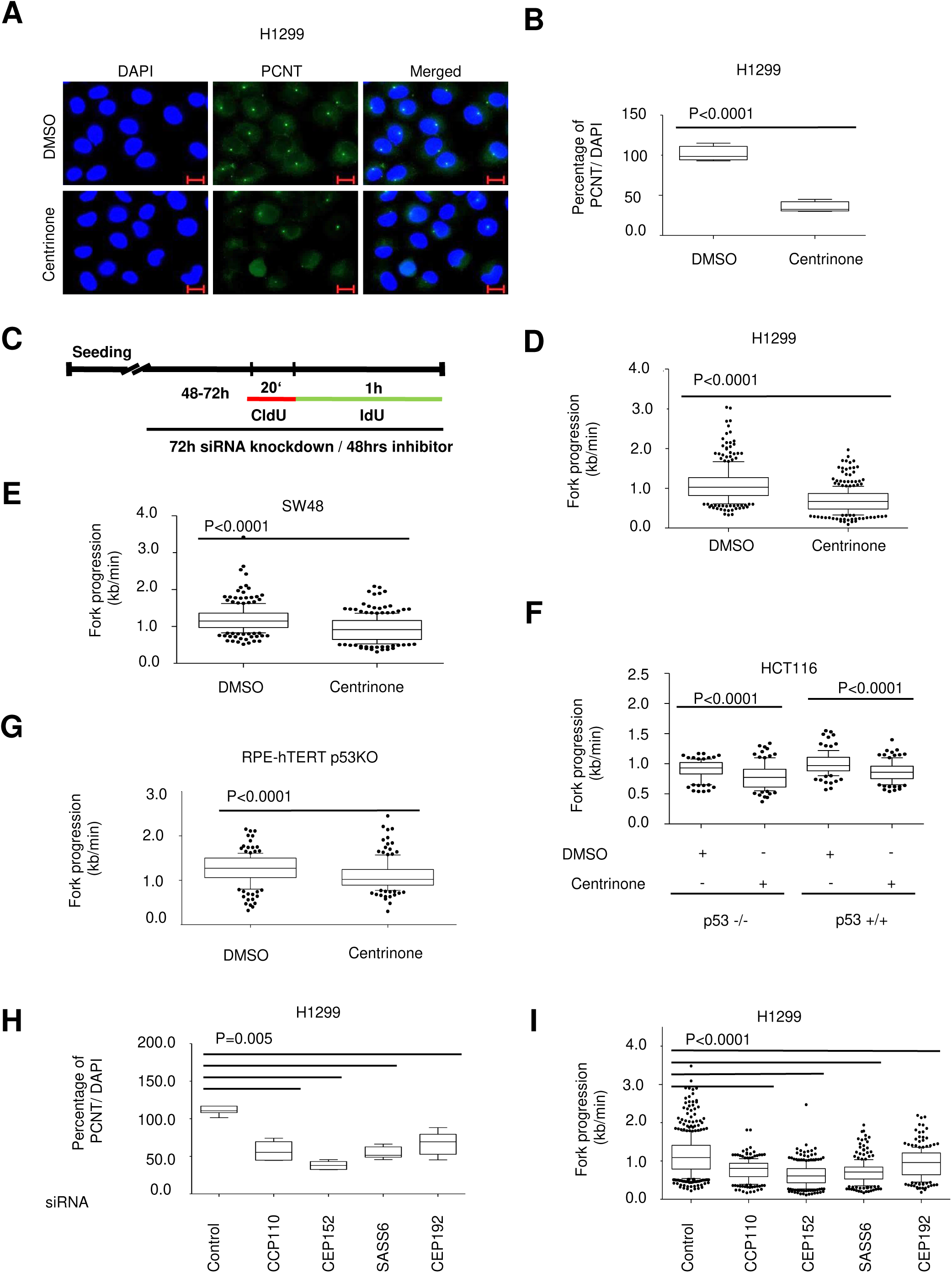
Depletion of centrosomal components interferes with DNA replication. **(A)** Detection of centrosome disintegration upon PLK4 inhibition. H1299 cells were treated with 500 nM Centrinone for 48 hrs. Centrosomes (immunostaining of PCNT) and the cell nuclei (DAPI) were detected (scale bar = 20 μm). Supplementary Figure 1 A, B represents the results of the same treatment, with staining of a different centrosomal component, CEP152. **(B)** Quantification of the centrosome signals per cell to in relation to DAPI-stained nuclei. Three replicates from A, with 150 cells each, were quantified per condition and presented as percentage (number of detectable centrosomes divided by the number of nuclei, multiplied by 100%) using GraphPad Prism. ****P < 0.0001. **(C)** Schematic workflow of cell treatment with Centrinone or pooled siRNA. H1299 cells were depleted of endogenous CEP152, CCP110, CEP192, or SASS6 by siRNA transfection for 72 hrs or treated with 500 nM Centrinone for 48 hrs and then labelled with CldU (20 min) and IdU (60 min), followed by fiber assay analysis. **(D)** Reduced DNA replication fork progression in response to PLK4 inhibition in H1299 cells, detected by DNA fiber assays. Cells were treated as described in (A) and (C), with 500nM Centrinone for 48 hrs, followed by incubation with 5’-chloro-2’-deoxy-uridine and then with iodo-deoxy-uridine. Tracks of newly synthesized DNA were visualized by immunostaining of CldU (red) and IdU (green). Fork progression was determined through the length of the IdU label (kb/min), assuming that 1 μm corresponds to 2.59 kb. 250 fibers were measured per condition per biological replicate and represented as a box plot. Biological replicates shown in Supplementary Figure 1 C, and the raw data are provided in Supplementary Table 2. **(E)** Decreased fork progression in the cell line SW48 (colon carcinoma) upon PLK4 inhibition. Cells were treated as in (B) followed by fiber assay. 200 fibers were measured per condition. Biological replicate in Supplementary Figure 1 E, raw data in Supplementary Table 2. **(F)** TP53 does not detectably influence fork progression in cells with impaired centrosomes. HCT116 cells that either retain TP53 or contain a targeted deletion of it were treated with 500nM Centrinone for 48 hrs and then subjected to fiber assay analysis. 150 fibers were measured per each condition. Biological replicates in Supplementary Figure 1 P, raw data in Supplementary Table 2. **(G)** Impact of PLK4 inhibition on non-transformed cells. Retinal pigment epithelial (RPE) cells immortalized with hTert and with a deletion of TP53 were treated as in (A) and (C), and DNA replication was analyzed by fiber assay. 150 fibers were measured per condition. Replicate in Supplementary Figure 1 J, raw data in Supplementary Table 2. **(H)** Quantification of the centrosome signals per cell to in relation to DAPI-stained nuclei. In three independent experiments, 300 cells in each were quantified per condition and presented as percentage (number of detectable centrosomes divided by the number of nuclei, multiplied by 100%). ****P < 0.0001. Representative images in Supplementary Figure 1 L. **(I)** Compromised DNA replication fork progression upon centrosome depletion, as determined by fiber assay. Cells were treated as in H and then subjected to fiber assay analysis. 200 fibers were measured per condition. Biological replicates are shown in Supplementary Figure 1 O, and the raw data are provided in Supplementary Table 2.

Upon PLK4 inhibition, we also found reduced proliferation of H1299 as well as HCT116 cells, but only after prolonged treatment for more than 2 days (Supplementary Figure 1 P-R). Similarly, the overall DNA synthesis rate per cell, as revealed by EdU incorporation and fluorescence staining, was only mildly impaired initially, with severe reduction only seen after three days (Supplementary Figure 1 S, T). Thus, the observed reduction in DNA replication fork progression cannot simply be ascribed to a general impairment of cell proliferation. Likewise, the distribution of DNA content (cell cycle distribution) and the number of chromosomes per cell remained largely the same after 48 hrs, whereas these parameters were found altered only after 7 days post treatment (Supplementary Figure 1 U-W).

Taken together, these observations suggest that the disintegration of centrosomes reduces the progression of DNA replication forks, independent of its long term impact on cell division.

### PLK4 inhibition or the depletion of centrosomal components each induces a DNA damage response

We hypothesized that the slow progression of DNA replication forks in response to the disintegration of centrosomes might be due to replication stress. To further investigate this possibility, we detected and quantified additional parameters indicative of a corresponding stress response, i.e. proteins that are phosphorylated by kinases of the DNA damage response.

First, we examined the accumulation of γH2AX as a marker of DNA damage in cells with centrosome impairment. Indeed, upon PLK4 inhibition by Centrinone, γH2AX levels markedly increased (Figure 2 A, B). Moreover, ATR and CHK1 were found phosphorylated (Figure 2 C). The depletion of centrosomal components led to similar effects (Figure 2 D-H). Thus, interfering with the integrity of centrosomes induces a typical DNA replication stress response.

**Figure 2.**
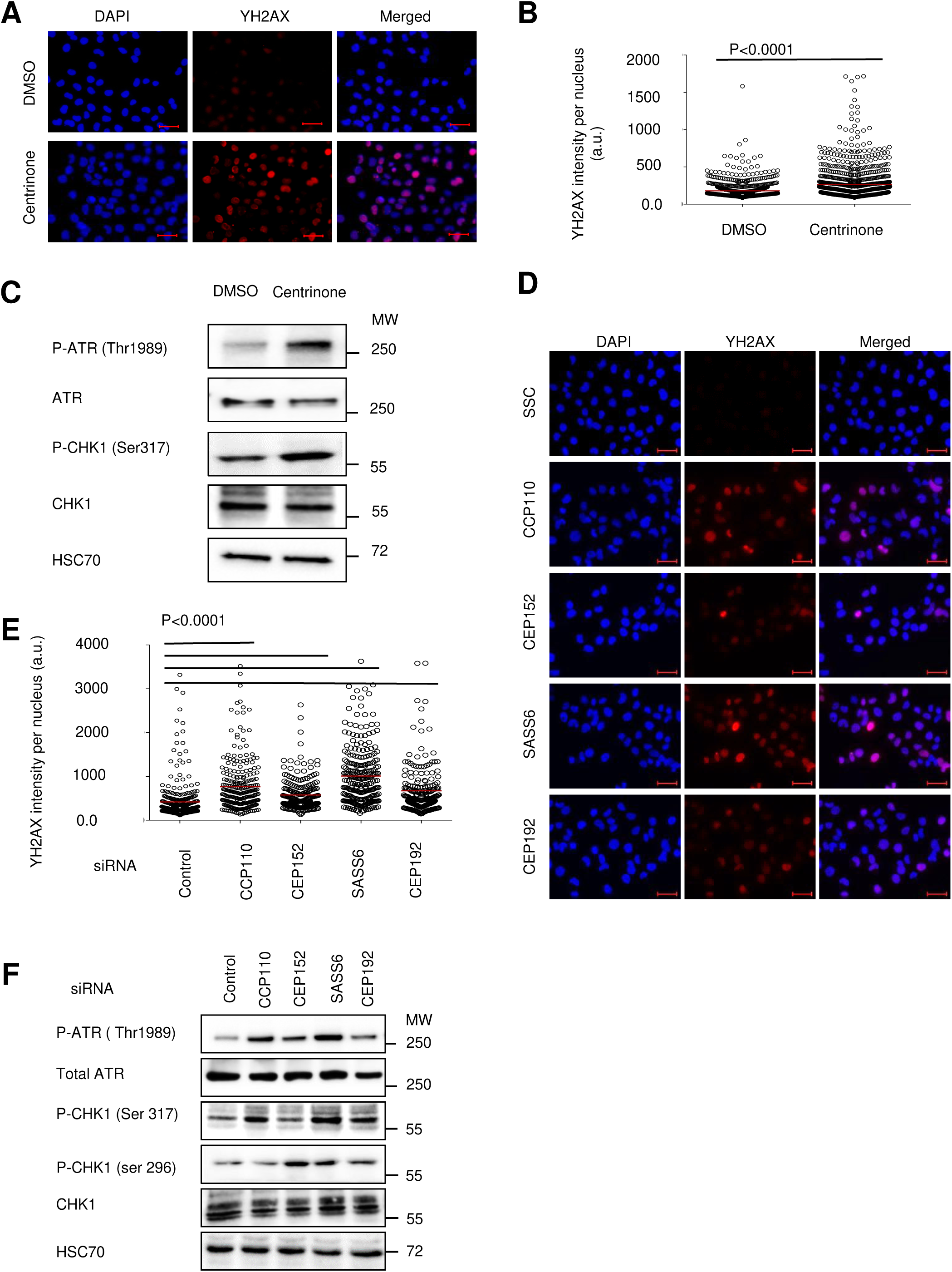
Centrosome impairment activates damage signaling. **(A)** Accumulation of γH2AX in cells treated with PLK4 inhibitor. Cells were treated with 500nM of Centrinone for 48 hrs, followed with immunostaining of Pericentrin (PCNT). Cell nuclei were counterstained with 4’,6-Diamidine-2’-phenylindole (DAPI). Scale bar, 20 μm. **(B)** Quantification of the γH2AX signal (A) using the ImageJ software. The mean (red bar) and distribution of three biological replicates (integrated) were calculated, and the level of significance was assessed by Mann-Whitney test. **(C)** Activation of ATR and CHK1 upon PLK4 inhibition. H1299 cells were treated with 500 nM Centrinone for 48 hrs and then subjected to immunoblot analysis to detect the indicated phosphorylations on ATR and CHK1. **(D)** Detection of γH2AX after depleting several centrosomal components. Cells were transfected with the indicated pools of siRNA for 72 hrs, followed with immunostaining as in (A). Scale bar, 20 μm. **(E)** Quantification of the γH2AX signal from (D) in cell nuclei upon centrosome depletion, as in (A). **(F)** Activation of ATR and CHK1 upon depletion of centrosomal components. Lysates of H1299 cells were prepared 72 hrs after transfection with pooled siRNAs as in (D), followed by immunoblot analysis.

### Impairment of centrosomes causes replication stress independent of mitosis

These results suggested to us that centrosomal integrity is required to maintain the processivity of DNA replication. However, it was not clear yet whether this effect is a direct one, or whether centrosome disruption first impairs chromosome segregation during mitosis, which might then impair DNA replication during the next S phase. The latter scenario was plausible for two reasons. Firstly, centrosome disruption often impairs the function of the mitotic spindle and thus chromosome segregation (Meraldi et al., 2016). Moreover, even one additional chromosome (numerical aneuploidy) is sufficient to trigger DNA replication stress (Passerini et al., 2016). We therefore developed a strategy of disrupting centrosomes and assessing DNA replication without allowing the cells to go through mitosis during the time of centrosome impairment. The technical difficulty in doing so consisted in the prolongation of the period of time required to deplete centrosomal components – a minimum of 72 hrs for siRNA knockdown or 48 hrs for PLK4 inhibition. Therefore, we sought to arrest the cells in G1 for 48 hrs and to disrupt the centrosome during this time. Only thereafter, the cells were released to S phase but not allowed to reach mitosis. To do so, we first arrested the cells in G1, using the cyclin dependent kinase 4 (CDK4) inhibitor Palbociclib (O’Leary et al., 2016). As shown in (Figure 3 A-B), this was achieved in less than 24 hrs. Washing off Palbociclib made the cells re-enter the cell cycle, but with variable time frames required for entering S (data not shown). To synchronize this entry, we released the cells from the CDK4 inhibitor but at the same time added thymidine, which is known to block the cell cycle right after entry into S (Whitfield et al., 2002). We then released the cells from the thymidine block for three hrs and thereby synchronized cells in S phase (Figure 3 A-B). In this way, we were now able to disrupt the composition of centrosomes and analyze DNA replication without intermediate mitosis. Control experiments revealed that the preceding CDK4 inhibition did not reduce the number of centrosomes, as had been found in different cell species (Adon et al., 2010) (Supplementary Figure 3 A, B). Using this system, we observed a substantial decrease in the progression of DNA replication forks again (Figure 3 C). Also, the overall DNA synthesis in the PLK4-inhibitor-treated cells was diminished after they had been released from the thymidine block (Figure 3 D, Supplementary Figure 3 D). Since these cells had not undergone mitosis during the time of PLK4 inhibition, the impairment of DNA replication cannot be ascribed to mitotic defects. Similar observations were made in synchronized cells when using the PLK4 inhibitor CFI-9400945 rather than Centrinone (Figure 3 C, E and Supplementary Figure 3 C, E), as well in PLK4-depleted cells (Figure 3 F and Supplementary Figure 3 F, G). Likewise, the depletion of centrosomal components led to a significant reduction in fork progression (Figure 3 G and Supplementary Figure 3 H), while the synchronization protocol was confirmed by cytometry in combination with knockdowns (Supplementary Figure 3 I). Thus, the disruption of the centrosomal composition interferes with the processivity of DNA replication, independent of chromosome missegregation.

**Figure 3.**
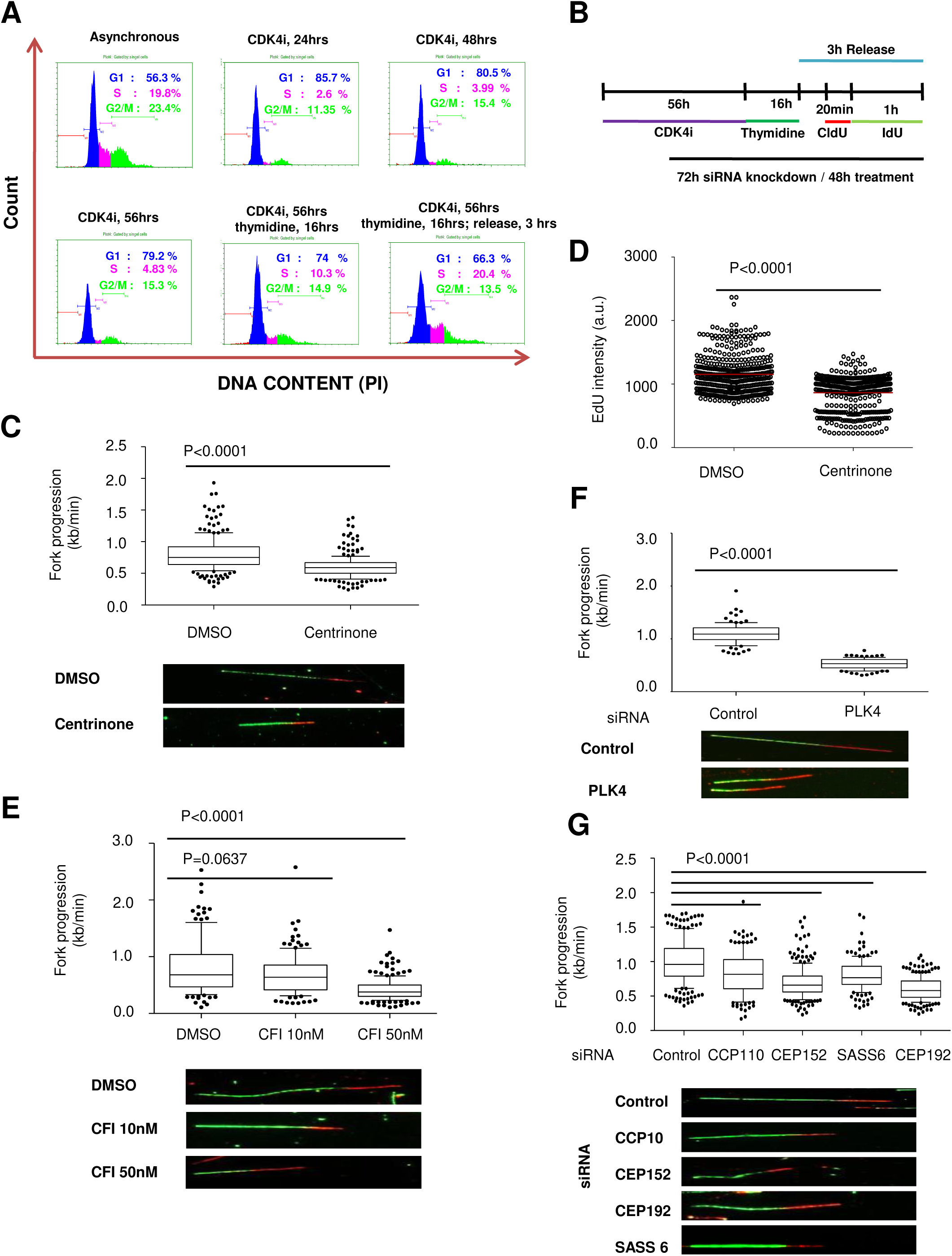
Impairment of centrosomes causes replication stress independent of mitosis. **(A)** Prolonged G1 arrest and synchronized entry to S phase. H1299 cells were treated with 5 μM CDK4 inhibitor (PD 0332991, also known as Palbociclib) for 24, 48 or 56 hrs. Cells treated with CDK4 inhibitor for 56 hrs were subsequently incubated with 2 mM thymidine for 16 hrs. Afterwards, the cells were washed and released into S phase for 3 hrs. DNA content and thus cell cycle distribution were assessed by propidium iodide (PI) staining and flow cytometry. **(B)** Schematic workflow of cell synchronization with CDK4 inhibitor (5 μM) and thymidine (2 mM), followed by labeling of newly synthesized DNA for fiber assay analysis. **(C)** Cells were synchronized as outlined in (B) and incubated with 5’-chloro-2’-deoxy-uridine followed by iodo-deoxy-uridine as indicated. Tracks of newly synthesized DNA were visualized by immunostaining of CldU (red) and IdU (green). Fork progression was determined through the length of the IdU label (kb/min). 200 fibers were measured per condition and represented as a boxplot. PLK4 was inhibited throughout the experiment (including the initial G1 arrest phase) by Centrinone (500 nM). A biological replicate is presented in Supplementary Figure 3 C. **(D)** Quantification of global DNA synthesis by 5-ethynyl-2’-deoxy-uridine (EdU) incorporation. Synchronized H1299 cells were treated with 500nM Centrinone for the indicated periods of time. Two hrs prior to fixation, the cells were incubated with 10µM EdU. After fixation, the incorporated EdU was visualized by click reaction between the alkyne moiety of EdU and a fluorescent Alexa dye coupled to azide, followed by quantification of the signal from individual cell nuclei (ImageJ; mean, red bar; distribution of 250 nuclei per condition). Representative images are shown in Supplementary Figure 3 D. **(E)** Cells were analyzed by fiber assays as in (C) after 48 hrs of treatment with another PLK4 inhibitor, CFI-400945, at the indicated concentrations. Biological replicate in Supplementary Figure 3 E. **(F)** Analyses of DNA replication fork progression as in (C) and (E). Rather than treating the cells with inhibitors, PLK4 was depleted using siRNA in H1299 cells. Knockdown efficiency and biological replicate presented in Supplementary Figure 3 F, G. **(G)** Fork progression was analyzed as in (C), after depleting the indicated centrosomal components with siRNA for 72 hrs. Biological replicate in Supplementary Figure 3 H.

### Centrosomal disintegration induces p38/MK2 signaling, and this is required for replication stress

Searching for the mechanisms that impair DNA replication fork progression upon centrosomal impairment, we tested the activity of p38/MK2 signaling, a pathway that we had previously found required for reducing DNA replication by nucleoside analogues or CHK1 inhibition (Kopper et al., 2013). Indeed, the phosphorylated and thus active forms of p38 and MK2 were strongly augmented by PLK4 inhibition (Figure 4 A), and the same was found for the bona fide MK2 substrate Hsp27 (Zheng et al., 2006). This is in agreement with previous reports on p38 activity affected by centrosomes (Mikule et al., 2007). Thus, the disruption of centrosomes activates p38/MK2 signaling, independent of mitotic dysfunction. Next, we tested whether the activation of p38/MK2 signaling is a cause of impaired DNA replication upon centrosome disintegration. We treated H1299 cells with the PLK4 inhibitor Centrinone. While assessing DNA replication using fiber assays, we incubated the cells with a pharmacological inhibitor of MK2 (Anderson et al., 2007). And indeed, DNA replication was restored to normal levels by interfering with MK2 activity (Figure 4 B, Supplementary Figure 4 A). Correspondingly, the levels of γH2AX were reduced, and cell proliferation was partially restored by MK2 inhibition in cells treated with Centrinone (Figure 4 C, D and Supplementary Figure 4 B). However, MK2 inhibition had no influence on centrosome numbers, neither in the presence nor in the absence of PLK4 inhibitors (Figure 4 E, Supplementary Figure 4 C), despite the reported ability of MK2 to phosphorylate the centrosomal protein CEP131 (Tollenaere et al., 2015). Moreover, we performed parallel experiments upon depletion of centrosomal components and co-depletion of MK2, with analogous results (Figure 4 F, Supplementary Figure 4 D). Similarly, MK2 activity was required to accumulate γH2AX and inhibit cell proliferation in this context (Figure 4 G-I, Supplementary Figure 4 E), but MK2 had has no impact on the numbers of centrosomes (Figure 4 J, Supplementary Figure 4 F). Hence, MK2 is a key mediator of replication stress upon disintegration of centrosomes.

**Figure 4.**
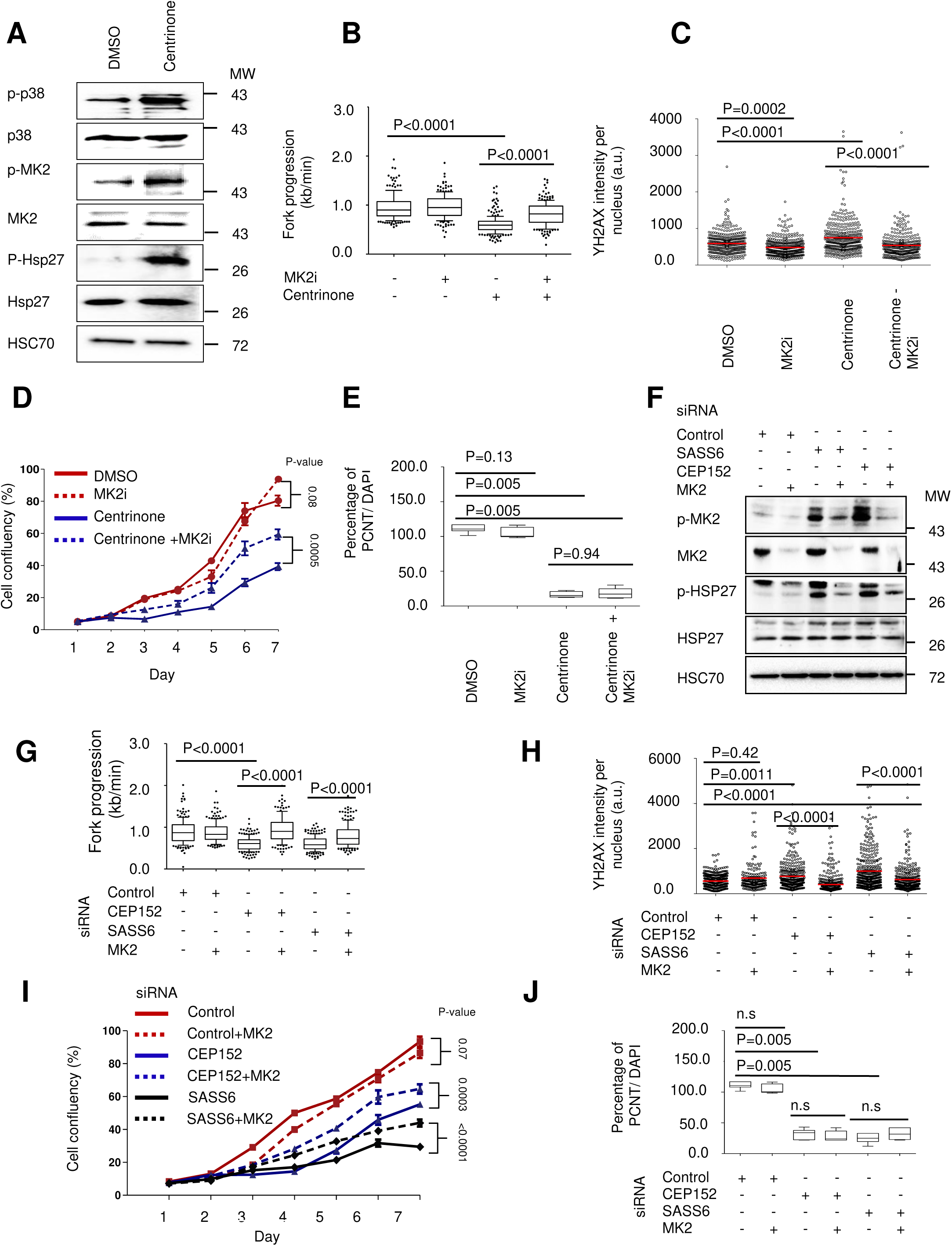
Centrosomal disintegration induces replication stress through p38 and MK2. **(A)** Activation of the p38/MK2 pathway by PLK4 inhibition. Synchronized (cf. Figure 3 B) H1299 cells were treated with 500 nM Centrinone for 48 hrs and analyzed by immunoblot. The phosphorylation of p38 and MK2, as well as the phosphorylation of the bona fide MK2 substrate HSP27, each indicate activation of the p38/MK2 signaling pathway. **(B)** Synchronized H1299 cells were treated with 500 nM Centrinone for 48 hrs, and with 10 μM of the MK2 inhibitor MK2iIII for the last 24 hrs, followed by fiber assays to quantitate DNA replication fork progression. Biological replicates shown in Supplementary Figure 4 A. **(C)** Quantification of the γH2AX signal in cell nuclei upon treatment with Centrinone and MK2iIII as in (B); images shown in Supplementary Figure 4 B. The nuclear signal was quantified using the ImageJ software. The mean and distribution of three biological replicates (integrated) were calculated, and the level of significance was assessed by Mann-Whitney test. **(D)** Partial rescue or cell proliferation by MK2 inhibition, in the context of PLK4 inhibition. 5*10^3^ H1299 cells were seeded in each well of a 24-well plate. Cells were treated with the DMSO solvent alone, or with 300 nM Centrinone, with or without 10 μM MK2iIII. Cell proliferation capacity was measured using the Celigo Cytometer (Nexcelom, software version 2.0). Confluence was measured every 24 hrs for 7 days. The experiment was carried out in three biological replicates with two technical replicates each. The mean and standard error were calculated from these six measurements at each time point. Note that most error bars are too narrow to be displayed. **(E)** MK2 activity has no impact on the number of centrosomes. Synchronized H1299 cells were treated as in (B) and then subjected to immunostaining of PCNT. 150 cells were counted per each condition, and an average of three biological replicates is represented in the figure. The number of detected centrosomes was divided by the corresponding number of DAPI-stained nuclei and displayed as a percentage. Representative images are shown in Supplementary Figure 4 C. **(F)** Activation of MK2 by depletion of centrosomal components. H1299 cells were reverse transfected for 72 hrs with 10 nM pooled siRNAs against CEP152 and SASS6. To assess the specificity of the pathway, MK2 was co-depleted in an additional set of samples. Immunoblot analyses revealed the phosphorylation of components in p38/MK2 signaling. **(G)** MK2 dependence of replication stress upon centrosome depletion. Synchronized H1299 cells were transfected as in F. Cells were subjected to fiber assays as in (B). Biological replicates shown in Supplementary Figure 4 D. **(H)** Quantification of the γH2AX signal in cell nuclei upon depletion of centrosomal components. H1299 cells were r transfected with 10 nM pooled siRNAs to knock down centrosomal components, in combination with siRNAs to MK2, for 72 hrs followed by immunostaining of γH2AX. Representative images are shown in Supplementary Figure 4 E. The nuclear signal was quantified using the ImageJ software. The mean and distribution of three biological replicates (integrated) are displayed. The significance was assessed by the Mann-Whitney test. **(I)** Partial rescue of cell proliferation upon centrosome depletion, by MK2 knockdown. H1299 cells were reverse transfected with 10 nM of siRNA as in (F), followed by assessment of cell proliferation as in (D) with three biological replicates / 6 technical replicates for each time point. **(J)** MK2 does not detectably affect the number of centrosomes. Synchronized H1299 cells were treated as in (F) and then subjected to immunostaining of centrosomes, using PCNT as a marker. 150 cells were counted per each condition, in three biological replicates. Calculations were performed as in (E), and representative images are provided in Supplementary Figure 4 F.

Conversely, overexpressing PLK4 not only enhanced the detectable centrosomal material but also enhanced fork progression and decreased MK2 signaling, even when cells were treated with gemcitabine, a nucleoside analogue that induces replication stress (Supplementary Figure 4 G-I). Thus, compared with PLK4 inhibition, enhanced PLK4 levels and accumulated centrosomal material have the reverse effects on DNA replication.

Taken together, the observed MK2 activation upon centrosome disintegration causes reduction in DNA replication fork progression, accumulation of γH2AX, and partially restored cell proliferation. Thus, MK2 may act as part of a surveillance pathway that ensures centrosome integrity. In response to centrosome impairment, MK2 interferes with DNA replication and cell proliferation, perhaps avoiding the accumulation of cells undergoing aberrant mitoses and chromosome missegregation.

### Upon centrosome disruption, the kinase MLK3 activates p38 and MK2

Given the crucial function of MK2 in replication stress, we sought to determine the upstream signaling pathway that leads to its activation in response to centrosome disruption. An upstream kinase of p38 that was previously found associated with the centrosome is MLK3 (Vertii et al., 2016). Accordingly, we found MLK3 associated with centrosomal structures, but only to a far lesser extent when the cells had been treated with the PLK4 inhibitor Centrinone (Figure 5 A, B). Strikingly, MLK3 inhibition prevented the accumulation of phosphorylated p38 and MK2, which otherwise occurred upon PLK4 inhibition (Figure 5 C), in agreement with an earlier report suggesting this possibility (Vertii et al., 2016). Moreover, MLK3 inhibition rescued DNA replication fork progression and abolished the accumulation of γH2AX in the presence of PLK4 inhibitor (Figure 5 D, E, Supplementary Figure 5 A, B). Interestingly, MLK3 inhibition partially restored the number of centrosomes in the context of PLK4 inhibition (Figure 5 F, Supplementary Figure 5 C). Similarly, MLK3 depletion largely restored DNA replication and decreased the levels of γH2AX when centrosomal components were knocked down (Figure 5 G-I, Supplementary Figure 5 D, E), and it prevented p38/MK2 activation (Figure 5 G). And again, knocking down MLK3 partially restored the number of detectable centrosomes when CEP152 was depleted (Figure 5 J, Supplementary Figure 5 F). Perhaps not surprisingly, no such rescue of centrosome numbers through MLK3 depletion was observed after knocking down the centriolar component SASS6. Thus, much like MK2, MLK3 is required for a signal triggered by centrosome disruption to interfere with DNA replication. In addition, however, MLK3 functions to further disintegrate the centrosome when PLK4 is inhibited or when a peripheral component of the centrosome, CEP152, is depleted. In any case, MLK3 activates p38 and MK2 in response to centrosome impairment, and this then causes replication stress.

**Figure 5.**
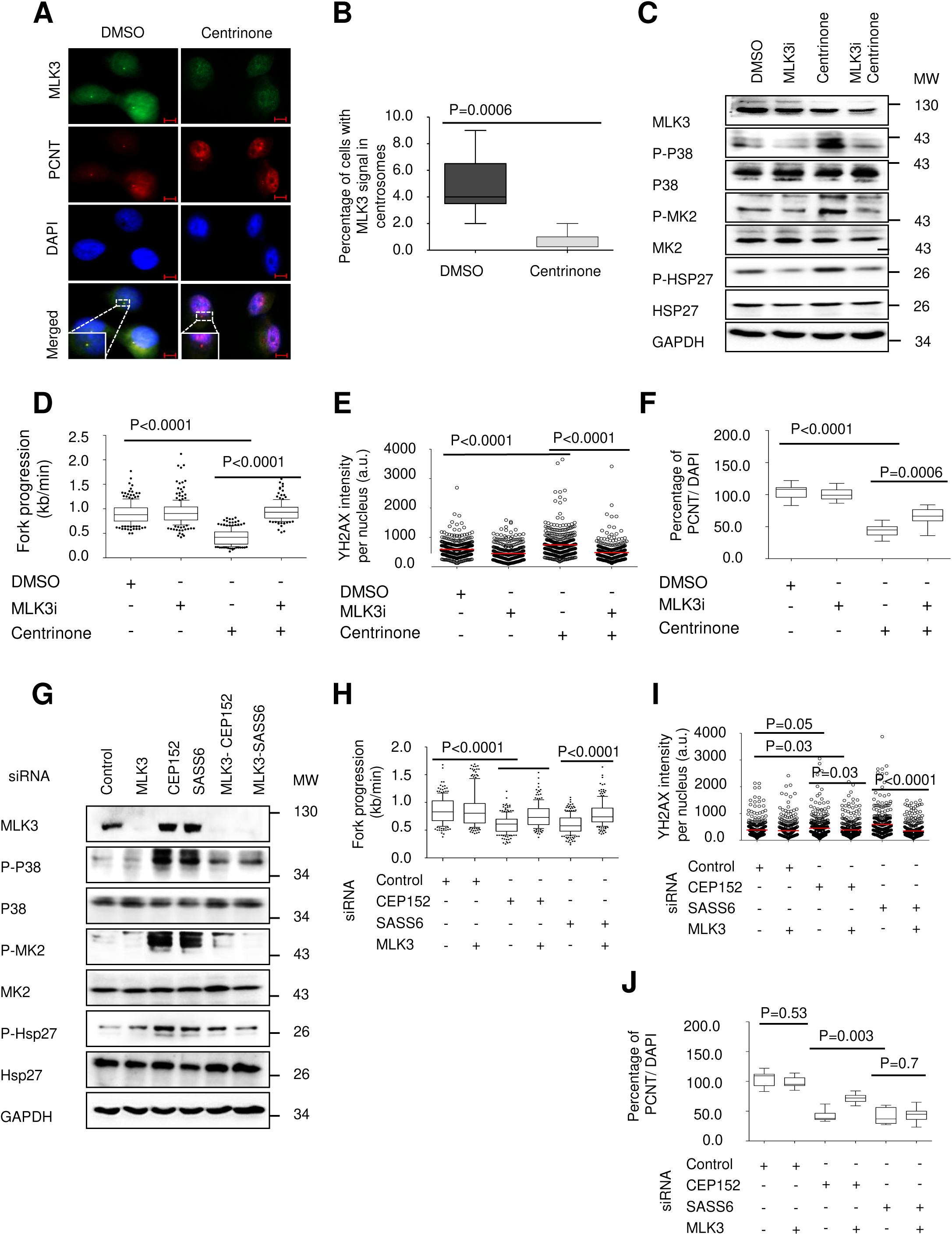
When centrosomes are disintegrated, MLK3 activates p38/MK2 to reduce fork progression. **(A)** MLK3 associates with centrosomes in a PLK4-dependent fashion. Synchronized H1299 cells were treated with either DMSO or 500 nM Centrinone for 48 hrs. Centrosomes were stained to detect PCNT and MLK3. DAPI was used to stain the nuclei (scale bar = 20 μm). **(B)** Quantification of MLK3 association to the centrosomal PCNT signal. 300 nuclei were quantified per condition. The results represent the average of three biological replicates with two technical replicates each. Some cells that were treated with Centrinone did not lose the centrosome signal (PCNT), and these were the cells that were included in the analyses regarding the colocalization of centrosomes with MLK3. ****P < 0.0001. **(C)** Dependence of p38/MK2 activation on MLK3. Synchronized H1299 cells were treated with 500 nM Centrinone for 48 hrs. During the last 24 hrs, the MLK3 inhibitor URMC-099 was added at 200 nM, followed by immunoblot analysis of p38/MK2 signaling. **(D)** Rescue of DNA replication in PLK4-inhibitor-treated cells by MLK3 inhibition. Synchronized cells were treated as in (C). DNA fiber assays were performed to assess fork progression. Biological replicates are shown in Supplementary Figure 5 A. **(E)** Suppression of γH2AX by inhibiting MLK3, in the context of PLK4 inhibition. The γH2AX signal in cell nuclei was quantified upon treatment with Centrinone for 48 hrs, and with 200nM MLK3 inhibitor for the last 24 hrs (representative images in Supplementary Figure 5 B). The mean and distribution of three biological replicates (integrated) were calculated, and the level of significance was assessed by Mann-Whitney test. **(F)** MLK3 inhibition can partially restore the number of centrosomes upon PLK4 inhibition. Quantification of the centrosome signals per cell to in relation to DAPI-stained nuclei. Synchronized H1299 cells were treated with 500 nM Centrinone for 48 hrs and 200nM of the MLK3 inhibitor URMC-099 for the last 24 hrs. Triplicates with 150 cells each were quantified per condition and presented as percentage (number of detectable centrosomes divided by the number of nuclei, multiplied by 100%). ****P < 0.0001. Representative images are shown in Supplementary Figure 5 C. **(G)** MLK3 knockdown diminishes p38/MK2 activation upon centrosome depletion. Synchronized H1299 cells were reverse transfected with 10 nM siRNAs against the targets CEP152, SASS6 and MLK3, in the indicated combinations, for 72 hrs, and pathway activation was detected by immunoblot analysis using phospho-specific antibodies. **(H)** MLK3 depletion rescues DNA replication when centrosomal components are removed by siRNA. Synchronized cells were treated as in (G) and then subjected to fiber assays. Biological replicate in Supplementary Figure 5 D. **(I)** Quantification of the γH2AX signal in cell nuclei upon depletion of centrosomal components, with and without MLK3 knockdown (images in Supplementary Figure 5 E). 72 hrs after transfection, γH2AX was quantified as in (E). **(J)** Depletion of MLK3 partially rescues centrosome numbers when knocking down CEP152. Quantification of detectable centrosomes in relation to DAPI-stained nuclei. Synchronized H1299 cells were transfected as in (G), and centrosomes were quantified as in (F). Representative images in Supplementary Figure 5 F.

### Centrosome disintegration induces RNA:DNA hybrids, and this is required for replication stress

Replication stress is often driven by unscheduled transcription and the formation of R-loops, i. RNA hybridizing to DNA (often in association with transcription) and displacing the opposite DNA strand (Aguilera and Garcia-Muse, 2012). Accordingly, upon PLK4 inhibition, we detected the formation of RNA:DNA hybrids by immunostaining with the monoclonal antibody S9.6 directed against these structures (Figure 6 A, B). Upon staining fixed cells in situ, the immunofluorescence signal derived from antibody binding was prominent in discrete nuclear structures, compatible with the concept that R-loops mainly occur at specific sites of highly active transcription (Tsekrekou et al., 2017). In contrast, the overexpression of RNaseH1, an RNase that specifically cleaves the RNA component of RNA:DNA hybrids, strongly reduced the nuclear immunostaining signal, confirming the specificity of the antibody (Figure 6 A, B). Similarly, using dot blot analyses, the accumulation of R loops was observed upon PLK4 inhibition but also when centrosomal components were depleted (Figure 6 C-E, Supplementary Figure 6 A, B). Treating the samples with RNaseH1 strongly reduced this signal (Figure 6 A-D), confirming RNA:DNA hybrids as the source of the antibody signal. Interestingly, the inhibition of p38/MK2 signaling also diminished the formation of R-loops upon centrosomal impairment, consistent with the rescue of DNA replication by the same inhibitors (Figure 6 F, G and Supplementary Figure 6 C, D).

**Figure 6.**
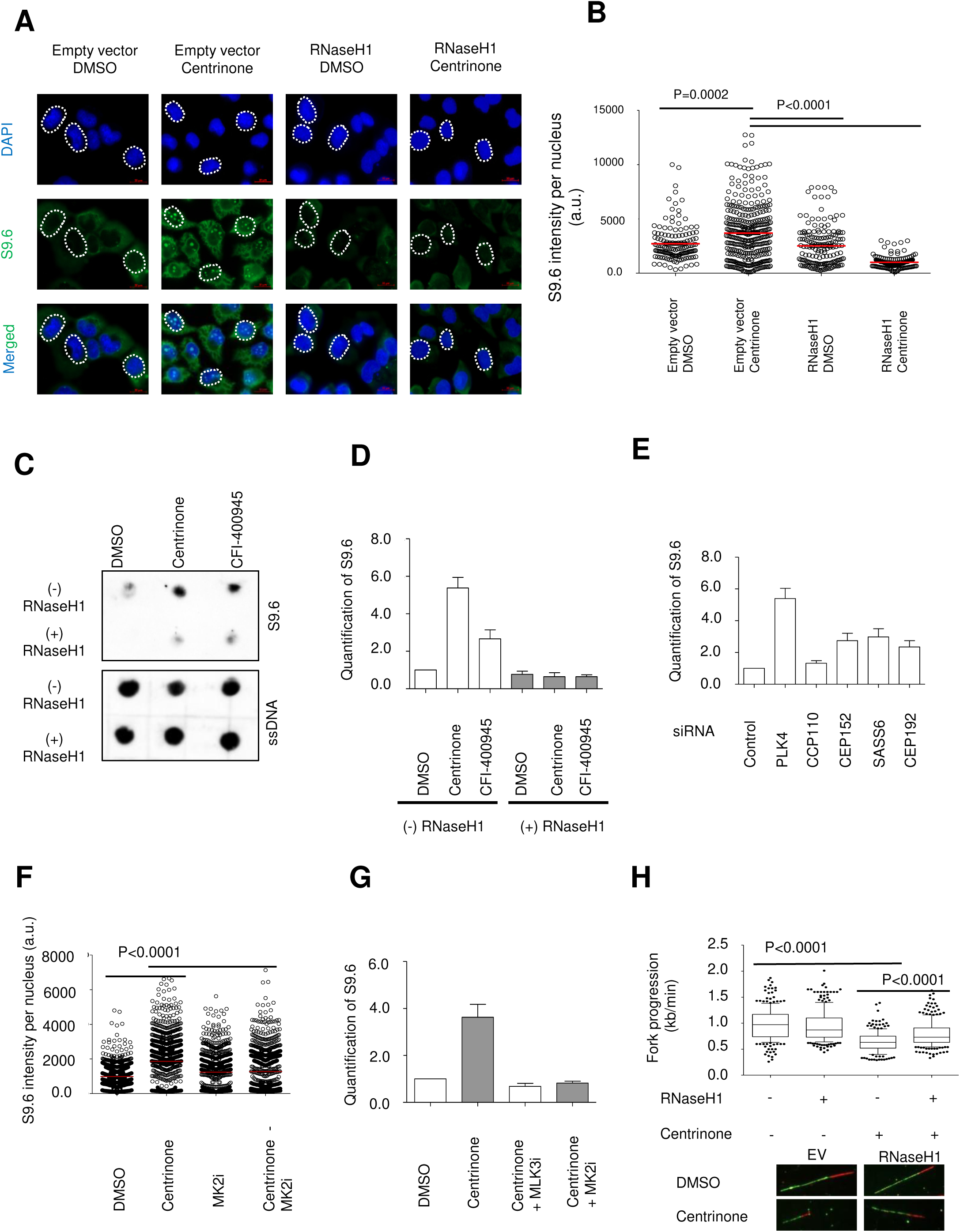
Replication stress upon centrosome disintegration requires RNA:DNA hybrids. **(A)** Immunostaining of the R-loop signal in cell nuclei upon treatment with Centrinone and/or transfection with an expression plasmid for RNaseH1. RNA:DNA hybrids were visualized by staining the cells with the antibody S9.6, and 4′, 6-Diamidin-2-phenylindol (DAPI) was used to delineate the nuclei (scale bar = 20 μm). **(B)** Quantification of the R-loop signal shown in (A). The nuclear signal was quantified using the ImageJ software. The mean and distribution of three biological replicates (integrated) were calculated, and the significance level was assessed using the Mann-Whitney test. **(C)** Dot-blot analysis of RNA:DNA hybrids in synchronized H1299 cells. Cells were treated either with DMSO, 500nM Centrinone or 10nM CFI-400945, for 48 hrs. The genomic DNA isolated from the cells was spotted onto a nitrocellulose membrane, with or without prior treatment with RNaseH1. The dots were stained with antibody S9.6 to detect RNA:DNA hybrids. For normalization, an antibody against single stranded DNA (ssDNA) was used to stain a parallel set of dots that had been acid-treated for denaturation. Biological replicates displayed in Supplementary Figure 6 A. **(D)** Quantification of the S9.6 signal on the membrane from (C). The signal obtained with S9.6 was normalized to the ssDNA signal first and then to the control treatment. Each column represents the average of three biological replicates with three technical replicates each. **(E)** Quantification of the S9.6 signal on the membrane as in (D), corresponding to the dot blots shown in Supplementary Figure 6 B. Calculations were carried out as in (D). A representative blot is shown in Supplementary Figure 6 B. **(F)** MK2 is required for R-loop formation upon PLK4 inhibition. RNA:DNA hybrids were detected by immunobluorescence upon treatment with Centrinone for 48 hrs, and with 10µM MK2iIII for the last 24 hrs. Quantification as in (B), representative images in Supplementary Figure 6 C. **(G)** MLK3 is required for R-loop formation upon PLK4 inhibition. Quantification of the S9.6 signal on the membrane corresponding to the dot blot in Supplementary Figure 6 D. The signal obtained with antibodey S9.6 was normalized to the ssDNA signal first and then to the control treatment. Each column represents the average of three biological replicates with three technical replicates each. **(H)** The removal of RNA:DNA hybrids by RNAseH1 rescues DNA replication fork progression upon Centrinone treatment. Cells were transfected with plasmids as in (A) and treated with Centrinone for 48 hrs. Fork progression was determined by fiber assays. Biological replicates are presented in Supplementary Figure 6 E.

To clarify the causal link between R-loop formation and replication stress, we performed fiber assays. Upon PLK4 inhibition, the progression of DNA replication forks was largely rescued by RNaseH1 overexpression (Figure 6 H, Supplementary Figure 6 E). We conclude that interfering with centrosomal integrity not only induces R-loop formation, but that this is a major cause of the observed DNA replication stress.

### PLK4 inhibition leads to the activation of the transcription factors, thereby enhancing replication stress

We next sought to determine mechanisms that may drive unscheduled transcription and lead to the formation of the RNA:DNA hybrids to induce replication stress, exploring potential transcription factors downstream of p38/MK2 signaling. Upon activation of p38 and MK2, the transcription factors ATF2 and JUN (also known as c-Jun) are phosphorylated and form a dimer to stimulate transcription (Breitwieser et al., 2007). Accordingly, we detected increased phosphorylation of ATF2 and JUN upon Centrinone treatment (Figure 7 A). At the same time, we observed the accumulation of both transcription factors in the chromatin fraction (Figure 7 B). ATF2/JUN phosphorylation was dependent on MK2 and MLK3 (Figure 7 C). JUN depletion restored DNA replication when PLK4 was inhibited (Figure 7 D, Supplementary Figure 7 A, B). Given the multitude of JUN-activated genes, this argues that JUN-mediated transcription might be partially responsible for the occurrence of replication stress. To investigate whether global transcription is indeed responsible for compromised DNA replication, we used an inhibitor of Cdk9 to shut down transcription, as described previously (Klusmann et al., 2016). And indeed, CDK9 inhibition rescued DNA replication in the presence of Centrinone (Figure 7 E, Supplementary Figure 7 C).

**Figure 7.**
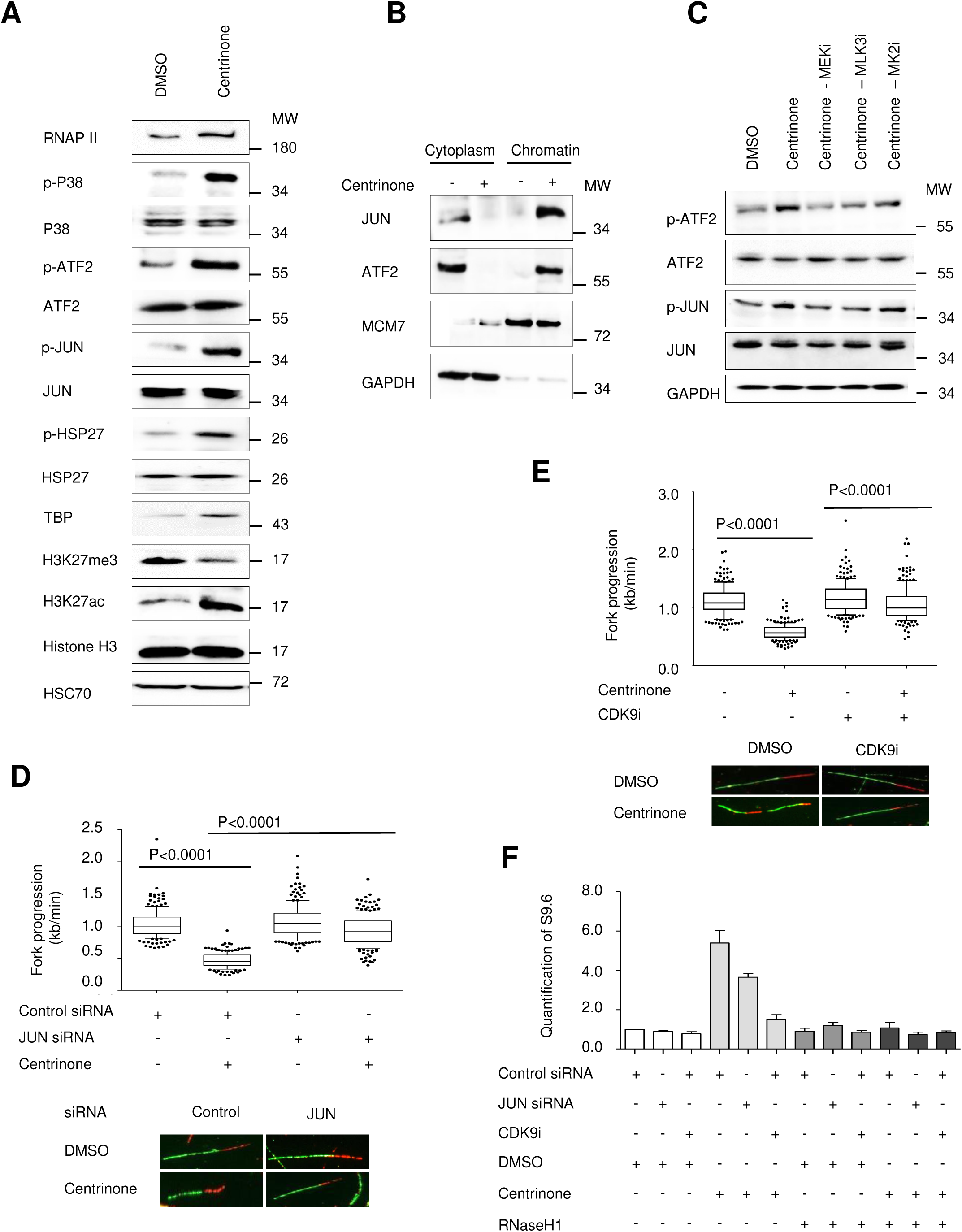
PLK4 inhibition activates ATF2 and JUN, and transcription is required to induce R-loops and replication stress. **(A)** Phosphorylation of ATF2 and JUN after Centrinone treatment (500 nM, 48 hrs), revealed by immunoblot analysis. Note the additional accumulation of TBP, the reduction in the repressive histone 3 trimethylation at K27, and the increase in H3K27 acetylation, all in agreement with increased transcriptional activity. **(B)** JUN and ATF2 associate with the chromatin fraction upon Centrinone treatment. Chromatin fractions were isolated and compared to the cytoplasmic supernatant. MCM7 (chromatin) and GAPDH (cytoplasm) were used to control the fractionation. The gel is a representative example of 3 biological replicates. **(C)** ATF2/JUN phosphorylation is dependent on the activities of MLK3 and MK2. H1299 cells were treated with the PLK4 inhibitor Centrinone as in (A), and in addition with MLK3i, MK2iIII and MEKi (U0126). The phosphorylation levels of ATF2 and JUN were assessed using immunoblot analysis. **(D)** The impairment of DNA replication fork progression upon Centrinone treatment is partially dependent on JUN. Synchronized H1299 cells were transfected to knock down JUN and treated with 500 nM Centrinone for 48 hrs. Fork progression was determined in fiber assays. Biological replicate in Supplementary Figure 7 A, B. **(E)** Blocking the global transcription machinery using a Cdk9 inhibitor rescues DNA fork progression upon PLK4 inhibition. Synchronized H1299 cells were treated with either DMSO or 500 nM Centrinone for 48 hrs, and 2 hrs before harvest the cells were additionally treated with 10µM Cdk9 inhibitor LDC000067. DNA fiber assays were performed to assess fork progression. Biological replicate in Supplementary Figure 7 C. **(F)** R-loop formation upon PLK4 inhibition depends partially on JUN and fully on global transcription. Quantification of the S9.6 signal from dot blot analyses (Supplementary Figure 7 D). Synchronized H1299 cells were treated with inhibitors and siRNAs as indicated underneath the figure. RNA:DNA hybrids. RNA:DNA hybrids were detected and quantified as in Figure 6 D. Each column represents the mean and SEM of three biological replicates with three technical replicates each.

To clarify the contributions of JUN and general transcription to R-loop formation, we performed dot blot analyses upon treatment of H1299 cells with the PLK4 inhibitor, along with JUN knockdown or CDK9 inhibition. This revealed that JUN depletion partially suppressed R-loop formation in response to PLK4 inhibiton. In contrast, blocking transcription with the CDK9 inhibitor fully reverted the accumulation of R-loops (Figure 7 F, Supplementary Figure 7 D). Thus, R-loop formation requires transcriptional activity, which is partially but not fully conferred by JUN activation upon centrosome disintegration.

Regarding replication stress, the rescue by JUN removal was more complete (Figure 7 D). We speculate that this might be due to the robustness of DNA replication. If R-loops need to accumulate above a certain threshold before DNA replication is impaired, this would explain why even a moderate reduction in R-loops, as observed upon JUN depletion, would still restore the progression of DNA replication forks.

### MK2 inhibition rescues defects in replication and proliferation of cells from patients with Seckel syndrome

The initial suspicion that centrosomes might govern DNA replication had come from the fact that genetic defects of centrosomes on the one hand, and the replication stress kinase ATR on the other hand, each lead to overlapping phenotypes in microcephaly (Kalay et al., 2011). We therefore asked whether the cells from patients suffering from a centrosomal defect might also display the features of replication stress. And indeed, human fibroblasts from a patient with a defect of the centrosomal component CEP152 (Kalay et al., 2011) showed substantially slower replication fork progression than the fibroblasts from healthy donors (Figure 8 A, Supplementary Figure 8 A). Moreover, the patient-derived cells had increased MK2 activity, as determined by the phosphorylation of HSP27, and they displayed enhanced levels of γH2AX (Figure 8 B). Strikingly, these cells also had increased amounts of p53 and its target gene product p21/CDKN1A, an inhibitor of cell cycle progression. MK2 inhibition markedly reduced the amount of detectable p21 in these cells, perhaps reflecting reduced p53 activity, albeit without affecting the levels of p53 itself (Figure 8 C). This is in agreement with the previously described ability of the p38/MK2 system to activate p53 through phosphorylation at Ser20 (She QB, 2002). Moreover, although patient-derived fibroblasts grew substantially less efficiently than normal fibroblasts, incubation with MK2 inhibitor led to equally efficient growth of fibroblasts from the patient or from healthy donors alike (Figure 8 D). Thus, centrosome disintegration in patients with a Seckel-related syndrome mediates MK2 activation and replication stress.

**Figure 8.**
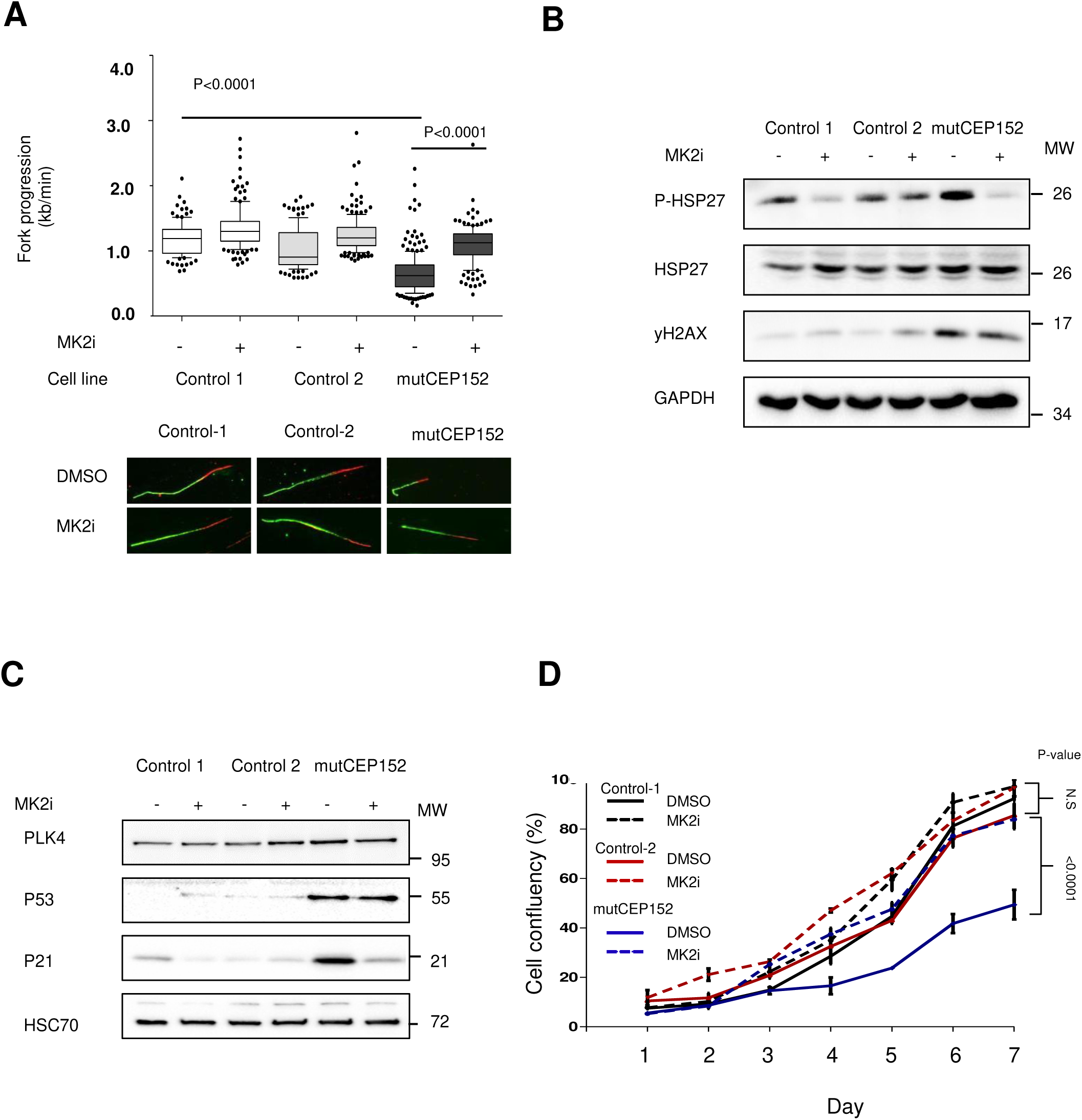
MK2 inhibition facilitates DNA replication and proliferation of cells derived from a Seckel patient. **(A)** The MK2 inhibitor MK2iIII rescues DNA replication fork progression in cells from a Seckel patient carrying a deletion in the CEP152 gene. Human breast fibroblasts (control-1), human skin fibroblasts (control-2) and skin fibroblasts from a patient carrying a homozygous mutation in CEP152 were treated with 20 µM MK2iIII for 48 hrs, followed by fiber assays. Data represented is an average of n=3. Biological replicate is shown in Supplementary Figure 8 A. **(B)** The MLK3-p38-MK2 pathway is constitutively activated in patient-derived cells with a mutation in CEP152. Cells were harvested after 48 hrs of 20 µM MK2iIII treatment and subjected to immunoblot analysis. **(C)** MK2 inhibition reduces the levels of p21/CDKN1A in patient-derived cells with a CEP152 mutation, despite the continuous accumulation of p53. The experiment was performed as in (B). **(D)** Restored proliferation rate of cells carrying a CEP152 mutation upon MK2 inhibition. 5*10^3^ fibroblasts from each patient were seeded in wells of a 24-well plate and treated with either DMSO or 10 μM MK2iIII. Cell proliferation was determined by a Celigo Cytometer. The average cell density from six replicates is presented at each time point.

## Discussion

Our results indicate that interfering with centrosomal integrity results in replication stress, even when cells are not allowed to undergo mitosis. Activation of MLK3-p38-MK2 signaling, through R-loop formation, mediates the reduction in DNA replication fork processivity. Thus, supporting unperturbed DNA replication represents a novel function of centrosomes.

A seemingly plausible mechanism of how interfering with the centrosomal function would result in replication stress is through malfunction of the mitotic spindle. When centrosomes disintegrate, mitotic fidelity can be decreased, thus enhancing the missegregation of sister chromatids. The resulting numerical chromosomal instability (CIN) would then result in replication stress, as was recently observed in cells with supernumerary chromosomes (Passerini et al., 2016). In continuously cycling cells, we do not rule out such a scenario. However, analysing chromosome spreads, we observed little change in ploidy during the treatment time with the PLK4 inhibitor, i.e. 48 hrs (Supplementary Figure 1 U-W). Moreover, by halting cell cycle progression during siRNA transfection or PLK4 inhibition, we show that centrosome disruption still leads to replication stress even without passage through mitosis (Figure 3). Rather than supernumerary chromosomes, the activation of MLK3-p38-MK2 signaling induces replication stress upon centrosome disintegration (Figures 4, 5).

The role of centrosomes in mitosis is subject to an ongoing debate. It became clear that centrosomes are not strictly required for the mitotic division of all cell types. Rather, they contribute to the accuracy of mitosis in some cell species but not others (Mikule et al., 2007). It is thus tempting to speculate that the role of centrosomes in DNA replication might be at least as important as their contribution to the accuracy of mitotic cell division. In terms of evolution, centrosomes may first have acquired their role as a microtubule organizing centre, maintaining the integrity of the cytoskeleton and of cilia. Such a role already requires precise timing of centrosome division. This, on the other hand, may have led to additional functions in the support of additional duplications during the cell cycle, i.e. the replication of DNA and the accuracy of the mitotic spindle.

We have previously characterized the role of MK2 in replication stress. In response to ultraviolet irradiation, treatment with the nucleoside analogue gemcitabine, or CHK1 inhibition, the resulting replication stress depends on MK2 and is relieved when MK2 is depleted or inhibited. At least in part, this rescue by MK2 inhibition is due to the reactivation of translesion synthesis polymerases, and accordingly, DNA polymerase eta was shown to represent an MK2 substrate (Kopper et al., 2013). We suggest that this signalling pathway is triggered by centrosome disruption as well, resulting in diminished progression of DNA synthesis when centrosomes are impaired.

The mixed-lineage kinase 3 (MLK3) was known to associate with centrosomes (Swenson et al., 2003) and to act upstream of MKK3/6 (Zhou et al., 2014). It is also known to activate JNKs and ERK, which may further induce replication stress (Chadee et al., 2006). Here we observed a functional interaction of MLK3 and the p38/MK2 complex. At least one of the functions of this interaction consists in the transmission of a signal that connects the replication of centrosomes and that of the cellular DNA.

Pharmacological inhibition of MLK3 using URMC099 was suggested for treatment of non-alcoholic steatohepatitis (NASH) (Tomita et al., 2017). Our results suggest that this drug may also re-enable DNA replication even under circumstances where MLK3 becomes active. It remains to be determined whether this might be beneficial by supporting liver regeneration, or whether it might support the outgrowth of cancer cell precursors by attenuating replication stress.

Initially, it was surprising to note that microcephaly associated syndromes often referred to as “Seckel syndrome” were found in patients with mutations of CEP152 (Kalay et al., 2011) or PCNT (Griffith et al., 2008), i.e. genes that encode centrosome components. The murine phenotype resembling human Seckel syndrome, including primary microcephaly, was also found in mice with a targeted deletion of CEP63, encoding another member of the centrosome (Marjanovic et al., 2015). Classically, both the human Seckel syndrome (O’Driscoll et al., 2003) and its murine model (Murga et al., 2009) were described in response to hypomorphic recessive alleles of ATR, the central mediator of the replication stress response which is required to dampen replication stress in most cells (Flynn and Zou, 2011). Mutations in the gene encoding the ATR-associated ATRIP protein also result in Seckel syndrome (Ogi et al., 2012). One way of explaining the similarity of syndromes consists in the concept that centrosomes might contribute to ATR signalling. Along this line, the ATR downstream kinase CHK1 was suggested to associate with centrosomes (Antonczak et al., 2016; Kramer et al., 2004). However, the association of proteins with centrosomes was suggested to be treated with caution due to numerous cross-reactions (Arquint et al., 2014). At least in our hands, CHK1 was not associated with centrosomes. Instead, our results strongly suggest that impaired centrosome composition triggers the translocation of MLK3, followed by activation of p38 and MK2. This signalling cascade induces replication stress, much like the deletion of ATR. This similarity in outcome makes it tempting to speculate that the disruption of ATR signalling, or the impairment of centrosome integrity by genetic alterations, can each lead to replication stress and thus to similar clinical conditions. However, we can currently not rule out that alternative mechanisms, e.g. the induction of p53 (Bazzi and Anderson, 2014; Fong et al., 2016; Lambrus et al., 2016; Meitinger et al., 2016), are more important determinants of microcephaly when centrosomes are defective. Moreover, it should be noted that syndromes that deviate from the classical Seckel phenotype can also occur in response to genetic alterations in centrosomal components (O’Neill et al., 2018). We consider the observed induction of replication stress upon centrosome disintegration as one but certainly not the only mechanism that might contribute to the clinical features when corresponding genes are impaired.

It is also somewhat surprising that the depletion of different centrosomal components results in similar patterns of replication stress. What these components do have in common is their requirement for centrosome duplication. Indeed, it has been reported previously that CCP110 (Chen et al., 2002), SASS6 (Clech et al., 2008), and CEP152 (Cizmecioglu et al., 2010) are each needed to duplicate the centrosome. Also, the inhibition of PLK4 led to aberrant centriole duplication during early stages of the cell cycle (Lei et al., 2018). Correspondingly, we observed reduced centrosome numbers upon depletion of each component. The failure to duplicate centrosomes might trigger the signaling chain reaching from MLK3 through p38 and MK2 to R-loops and replication stress. Another clue to the underlying mechanism is provided by the location of MLK3 on centrosomes, which is transient in nature and is temporarily interrupted shortly before mitotic entry (Swenson et al., 2003). This transient nature of the association at least suggests that MLK3 is not a constitutive part of the centrosome but represents a peripheral and quite lose association partner. As a consequence, we propose that multiple alterations in centrosome composition can dissociate MLK3 from the centrosome, leading to signalling activation in a uniform manner.

Taken together, the findings reported here suggest a novel function of centrosomes in the cell cycle. On top of providing microtubule organizing centres and a supportive function in chromosome segregation, centrosomes also enable the unperturbed progression of DNA replication. In addition to ATR/Chk1 signalling (Dobbelstein and Sorensen, 2015) and histone supply (Alabert et al., 2017), centrosomes support the progression of DNA replication forks.

## Acknowledgments

We thank Valentina Manzini for help in the computer-assisted quantitative image evaluation of immunofluorescence data. This work was supported by the Deutsche Forschungsgemeinschaft (FOR 2800), the Else Kröner Fresenius Stiftung, the Wilhelm Sander Stiftung, the Deutsche José Carreras Leukämie Stiftung, and the Deutsche Krebshilfe, ZT was a member of the Göttingen Graduate School GGNB during this work.

## Author contributions

MD conceived the project and drafted the manuscript. ZT, KS, AK, MF, JC and BW performed experiments. MD, ZT and BW revised the manuscript and all authors approved it.

## LEGENDS TO SUPPLEMENTARY FIGURES

**Supplementary Figure 1, corresponding to Figure 1.**
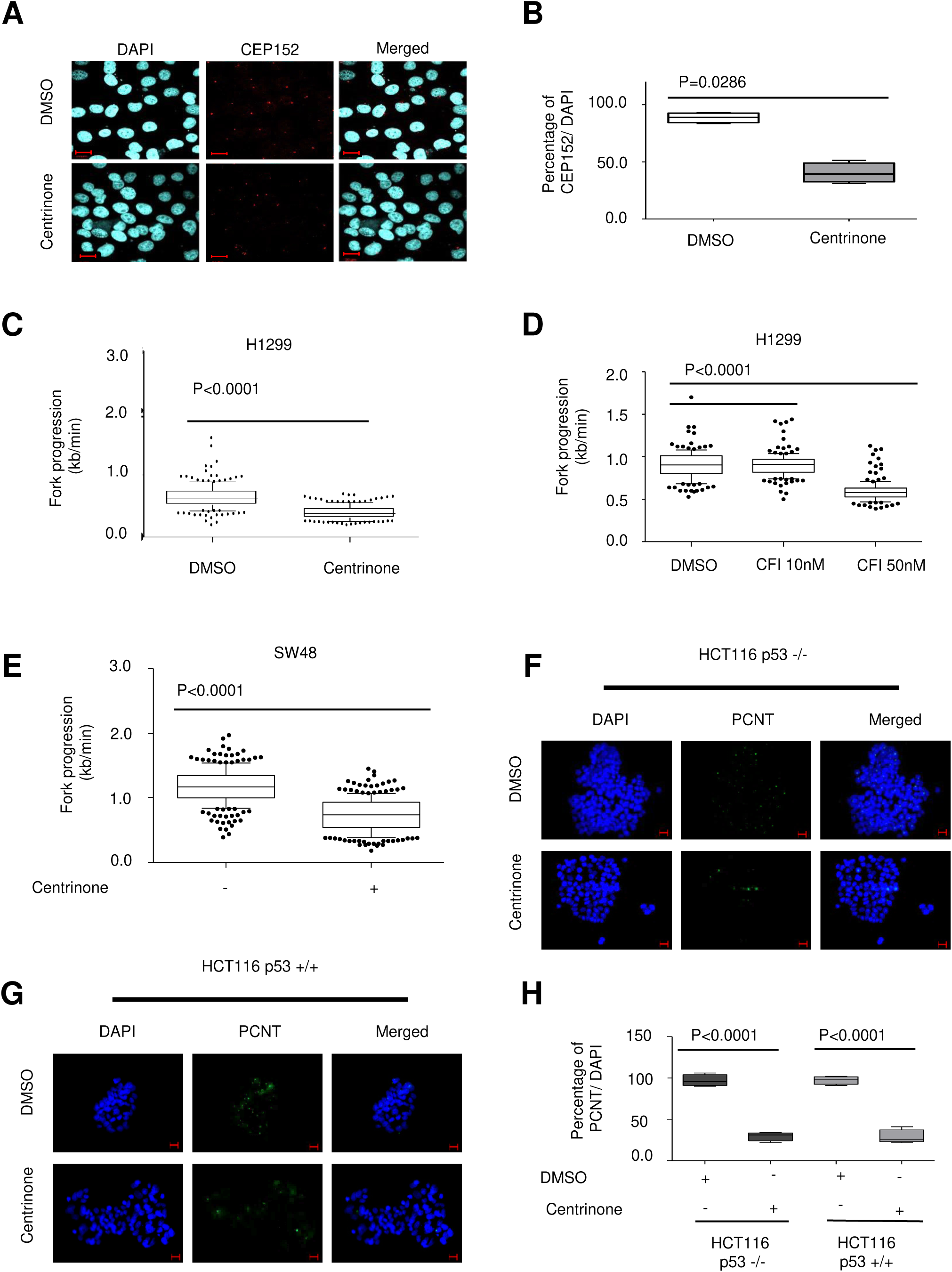
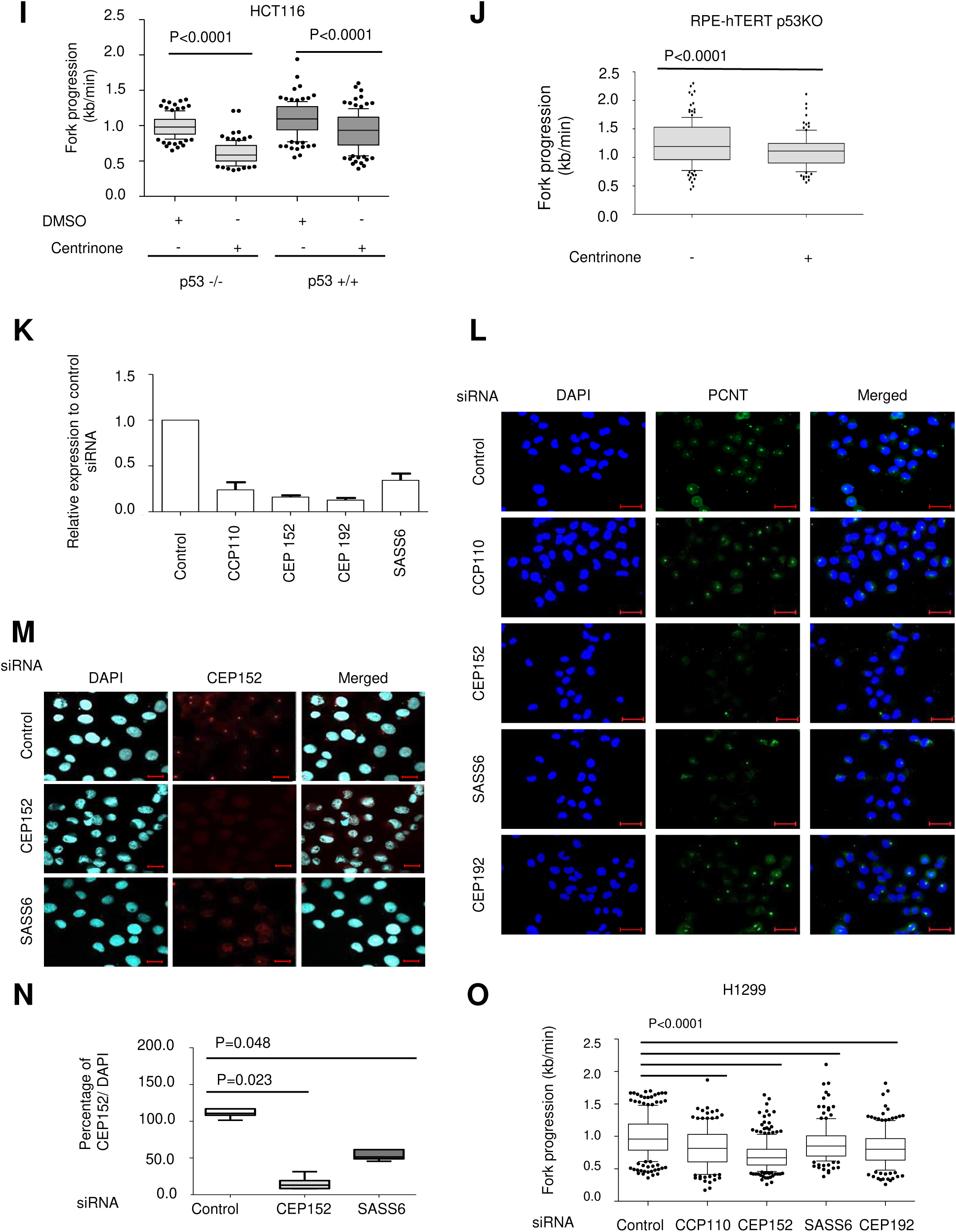
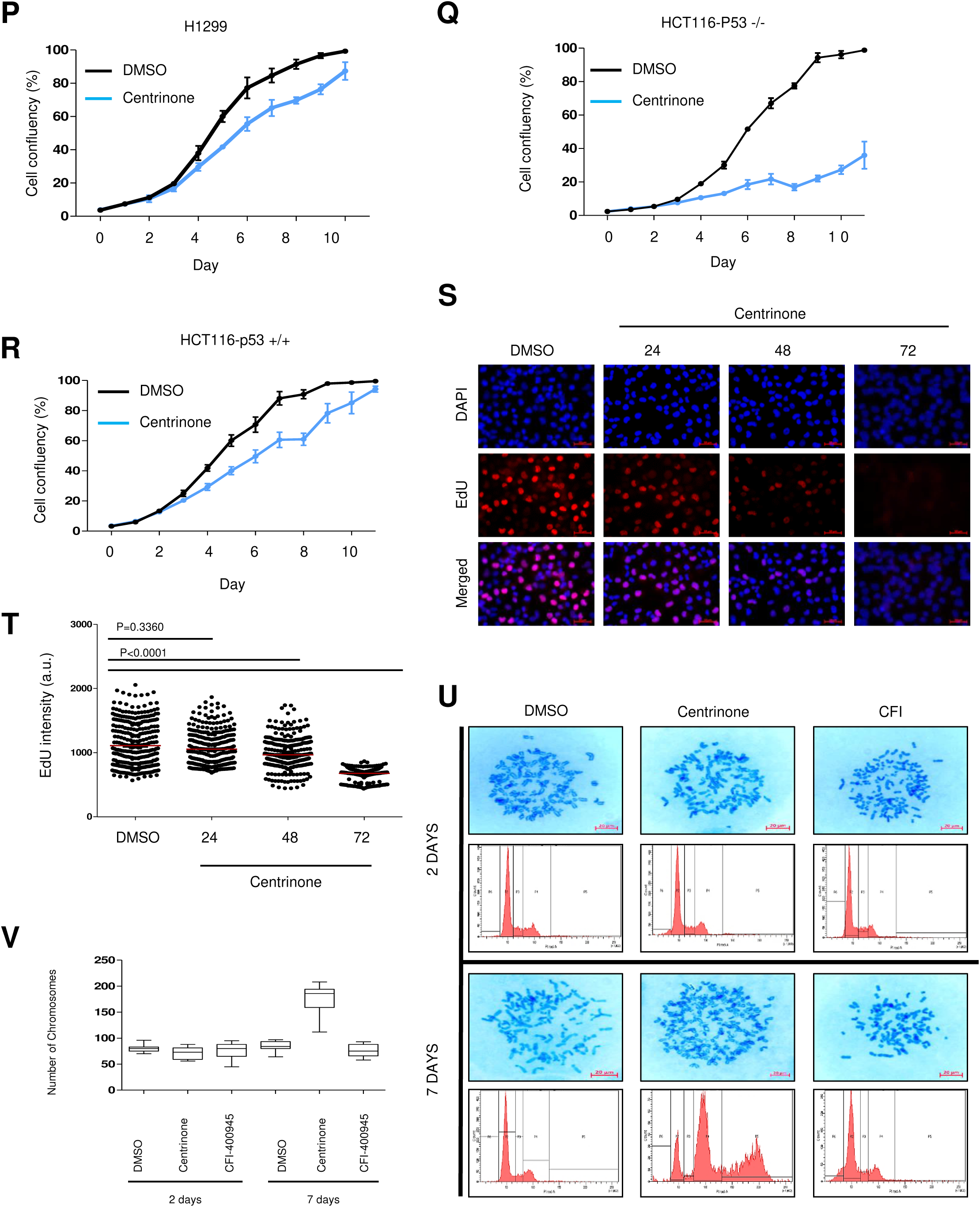
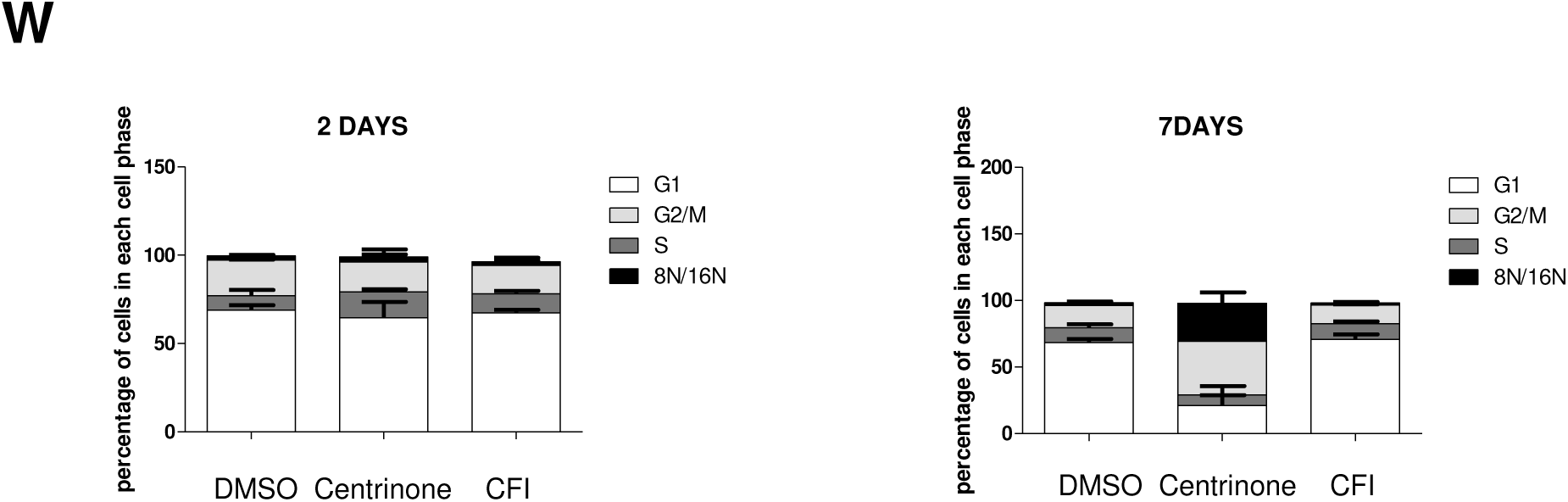
Centrosome disruption weakens the progression of DNA replication forks. Reduced DNA replication fork progression by DNA fiber assays in cells with the following properties. **(A)** Detection of centrosome disintegration upon PLK4 inhibition. Cells were treated with 500 nM Centrinone for 48 hrs. Centrosomes were detected by immunostaining of CEP152, and the cell nuclei were detected by 4′,6-Diamidin-2-phenylindol (DAPI; scale bar = 20 μm). **(B)** Quantification of centrosomes from A. Quantification of the centrosome signals per cell to in relation to DAPI-stained nuclei. 300 cells from A were quantified per condition and presented as percentage (number of detectable centrosomes divided by the number of nuclei, multiplied by 100%). ****P < 0.0001. **(C)** DNA replication fork progression, detected by fiber assays, in non-synchronized H1299 cells treated with 500 nM Centrinone, as in Figure 1D, and incubated with 5’-chloro-2’-deoxy-uridine followed by iodo-deoxy-uridine. Fork progression was determined through the length of the IdU label (kb/min). 200 fibers were measured per condition and presented as a boxplot. **(D)** Fiber assay on non-synchronized H1299 treated with PLK4 inhibitor CFI-400945 **(E)** Fiber assay on SW48 cells treated with 500 nM Centrinone, as in Figure 1 E. **(F)** Detection of centrosome disintegration upon PLK4 inhibition. HCT116 deficient in p53 were treated with 500 nM Centrinone for 48 hrs. Centrosomes (immunostaining of PCNT) and cell nuclei (DAPI) were detected (scale bar = 20 μm). **(G)** Detection of centrosome disintegration upon PLK4 inhibition in p53-proficient HCT116 cells as in (F). **(H)** Quantification of centrosomes from F and G. Quantification of the centrosome signals per cell to in relation to DAPI-stained nuclei. Three samples of 150 cells each were quantified per condition and presented as percentage (number of detectable centrosomes divided by the number of nuclei, multiplied by 100%) ****P < 0.0001. **(I)** Fiber assays performed using HCT116 cells (with or without p53) treated with Centrinone, as in Figure 1 F. **(J)** Fiber assays on RPE-hTert cells treated with Centrinone, as in Figure 1 G. **(K)** Knockdown efficiency for the depletion of centrosomal components. RNA was isolated from cells 48 hrs post siRNA transfection to knock down CCP110, CEP192, CEP152, and SASS6. Each mRNA was quantified by reverse transcription and quantitative polymerase chain reaction (qRT-PCR). The relative expression upon knockdown is displayed normalized to control siRNA #1, and to the house keeping gene HPRT1. Mean and SEM of n=3. **(L)** Representative images of centrosome staining upon knocking down the indicated components of centrosomes. The results are quantified in Figure 1H. **(M)** Centrosome disintegration upon depletion of centrosomal components. H1299 cells were transfected with pools of three siRNAs each, against CEP152 and SASS6, for 72 hrs. Centrosomes were detected by staining CEP152, and 4′,6-DAPI was used to delineate the nuclei (scale bar = 20 μm). **(N)** Quantification of the centrosome signals per cell to in relation to DAPI-stained nuclei as in (B). 300 cells from (M) were quantified per condition. ****P < 0.0001. **(O)** Fiber assays on H1299 cells in response to depleting several centrosome components, replicate to Figure 1I. **(P)** Reduction in cell proliferation upon PLK4 inhibition with Centrinone. 5*10^3^ H1299 cells were seeded in each well of a 24-well plate. Cells were treated with DMSO or 300 nM Centrinone. Cell proliferation capacity was measured using the CeligoTM Cytometer (Nexcelom, software version 2.0). Confluence was measured every 48 hrs for 10 days. The experiment was carried out in three biological replicates and 6 technical replicates for each time point. Note that most error bars are too narrow to be displayed. **(Q)** Proliferation of HCT116 cells with a targeted deletion of p53 upon PLK4 inhibition with Centrinone. The experiment was carried out as in (P). **(R)** Proliferation of p53-proficient HCT116 cells upon PLK4 inhibition, assayed as in (P) and (Q). **(S)** Overall DNA synthesis upon long term inhibition of PLK4. Representative images of the EdU incorporation signal in H1299 cells are shown. Cells were treated with Centrinone for 24-72 hrs, and EdU incorporation was visualized using a click reaction with an Alexa fluorescent dye coupled to azide. **(T)** Quantification of images from (S) using the ImageJ software. The mean and distribution of three biological replicates (integrated) were calculated. Significance, Mann-Whitney t-test. **(U)** Long term impact of PLK4 inhibition on cell ploidy. Upper panel: Representative images of chromosome spreads from three biological replicates. Cells were treated with 500nM Centrinone or 10nM CFI-400945 for two or seven days, followed by spreading and visualizing chromosomes. Chromosomes were stained with 8% Giemsa solution and images were acquired by microscopy (100x, bright field mode). Lower panel: cell cycle profiles corresponding to the chromosome spreading experiment. DNA content and thus cell cycle distribution were assessed by propidium iodide (PI) staining and flow cytometry. **(V)** The number of chromosomes per cell was counted manually from (U) and plotted using GraphPad Prism (n=3 cells per time point and condition). **(W)**Quantification of the cell cycle profiles from (U), average of three biological replicates.

**Supplementary Figure 2, corresponding to Figure 2**

There is no supplementary data to Figure 2.

**Supplementary Figure 3, corresponding to Figure 3.**
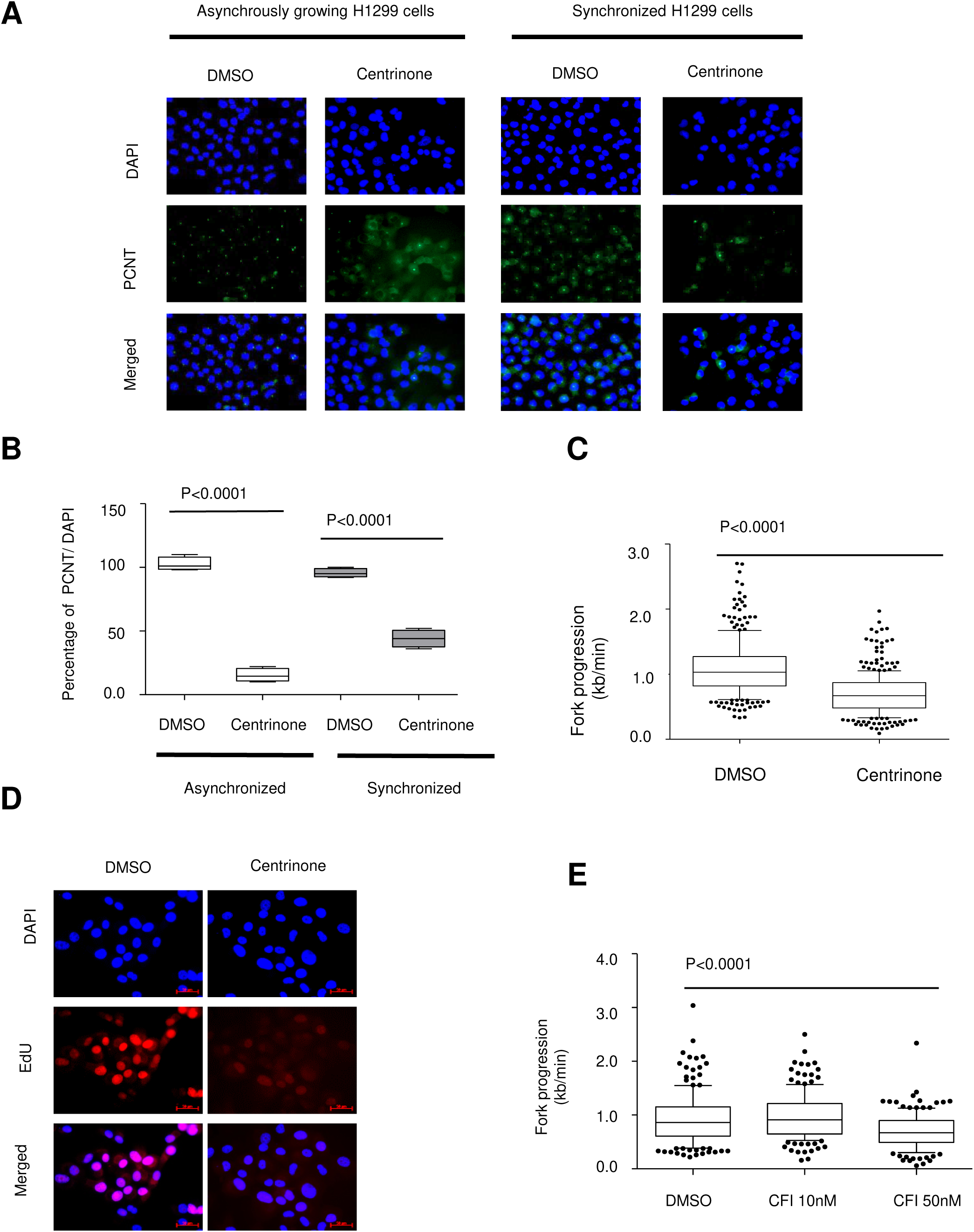
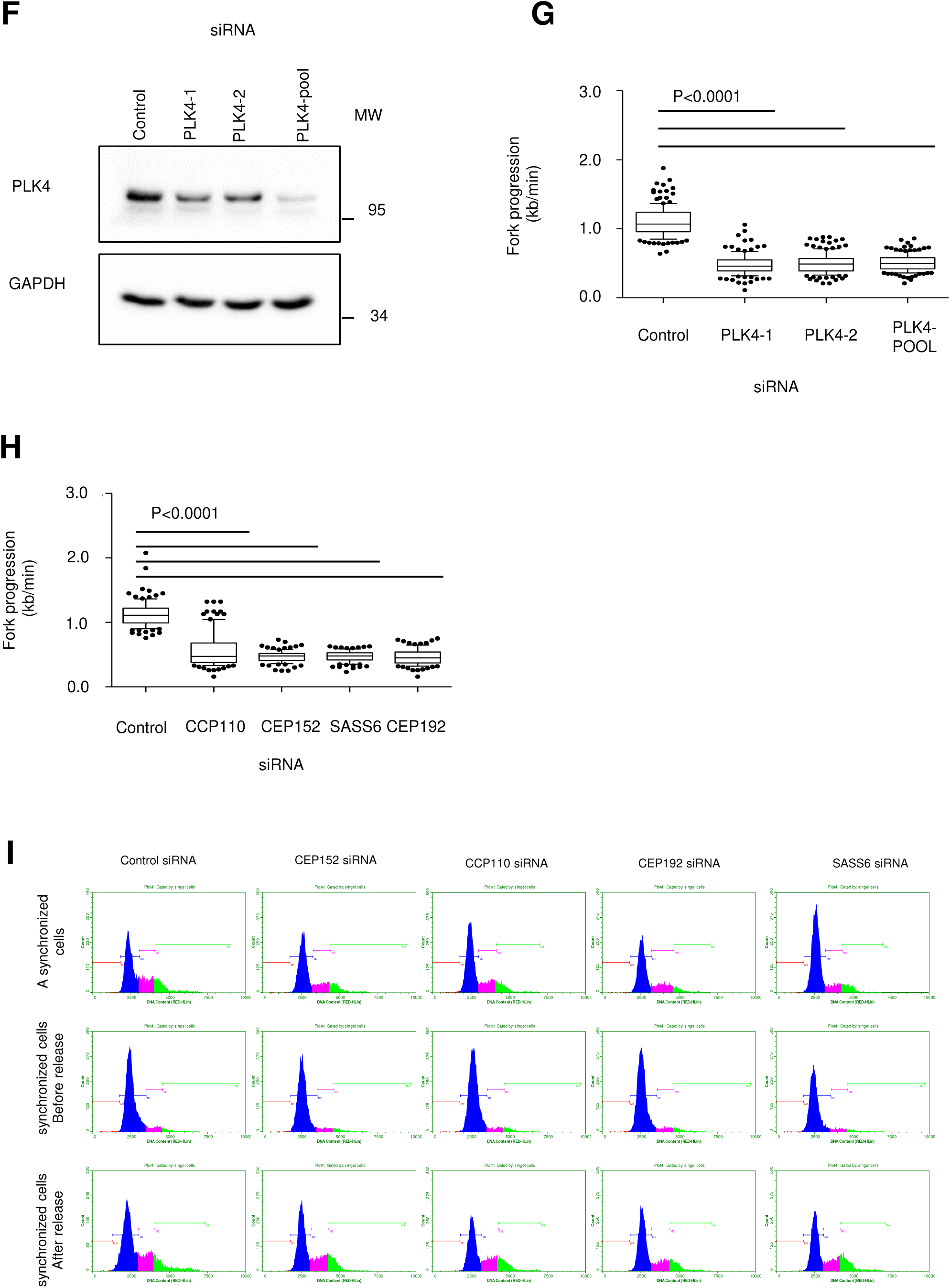
**(A)** Evaluating the effect of CDK4 inhibition on the numbers of centrosomes before and after synchronization. Cells were synchronized as in Figure 3 B, followed by immunostaining of PCNT as in Figure 2 A. **(B)** Quantification of centrosomes per cell nucleus, based on the data displayed in (A). **(C)** Replicate of the fiber assays on synchronized H1299 cells treated with Centrinone, as in Figure 2C **(D)** EdU incorporation of synchronized H1299 upon Centrinone treatment and release from the thymidine block. 2 hrs prior to fixation, the cells were incubated with 10µM EdU, followed by staining through click-reaction with an azide-coupled Alexa dye. Quantification provided in Figure 3 D. **(E)** Fiber assay, on synchronized H1299 cells upon treatment with the PLK4 inhibitor CFI-400945. Replicate to Figure 2E. **(F)** Validation for the PLK4 siRNA efficiency using immunoblot analysis. Single siRNAs and the pool of both were used. **(G)** Impaired DNA synthesis upon PLK4 depletion, assayed on synchronized H1299 cells treated as in Figure 2 F. **(H)** Depletion of centrosomal components resulting in diminished replication fork progression. Synchronized H1299 cells were treated as in Figure 2 G. **(I)** Distribution of DNA content (cell cycle distribution) of cells upon depletion of centrosomal component and synchronization. Flow cytometry was carried out upon siRNA transfection to deplete CEP152, CCP110, CEP192 or SASS6, on asynchronously growing cells, and on synchronized cells (the latter before and after release from a thymidine block).

**Supplementary Figure 4, corresponding to Figure 4.**
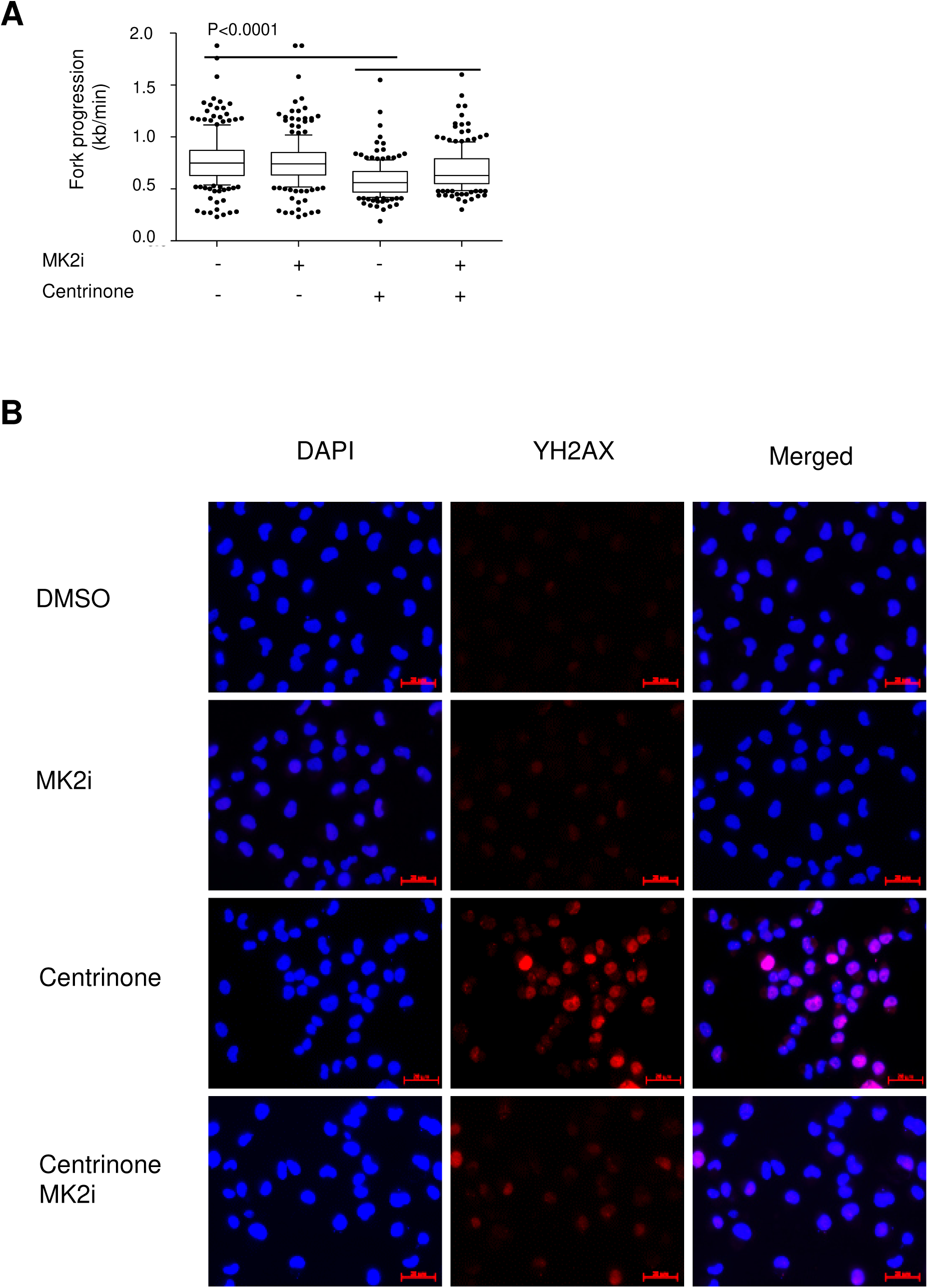
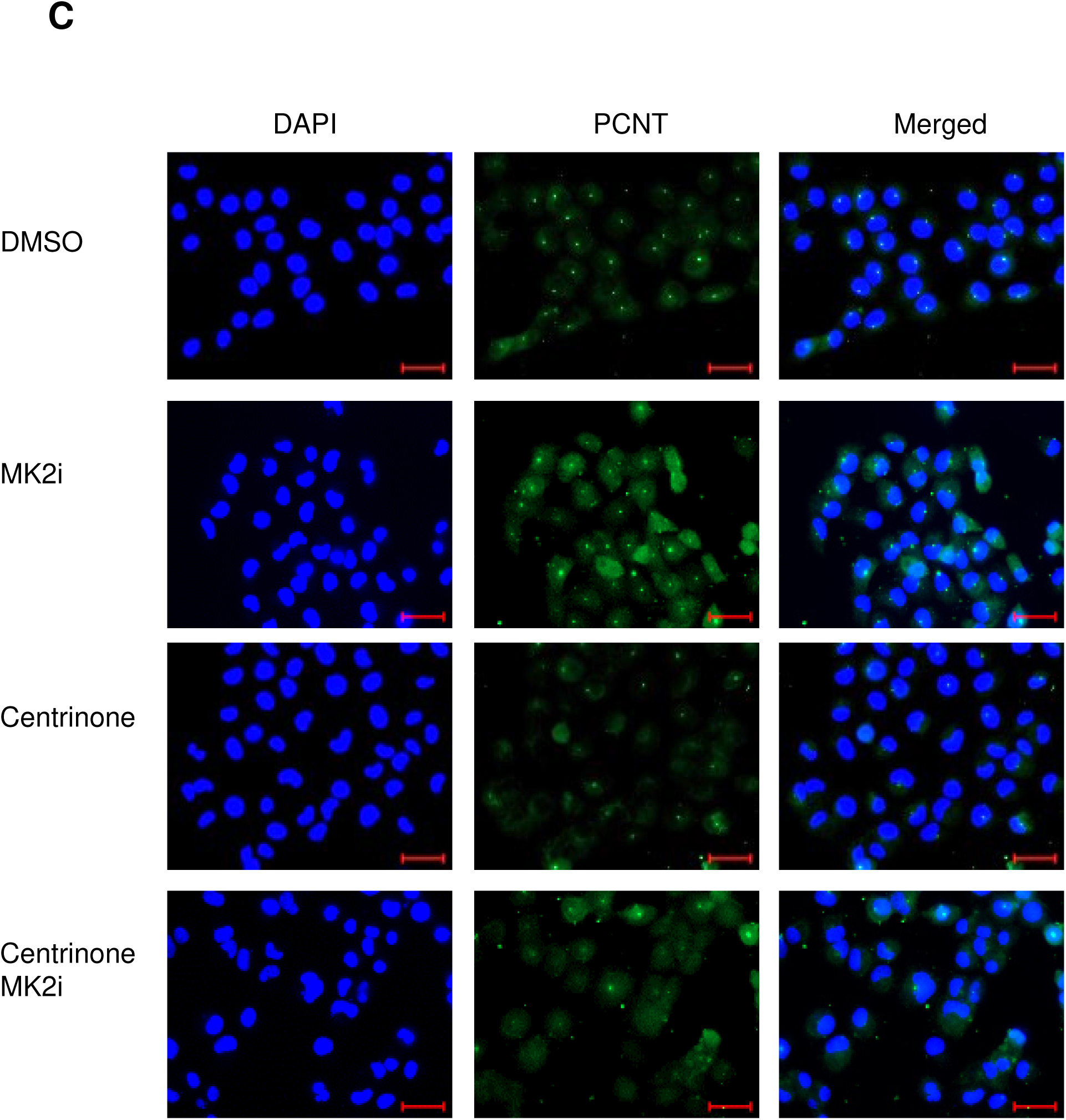
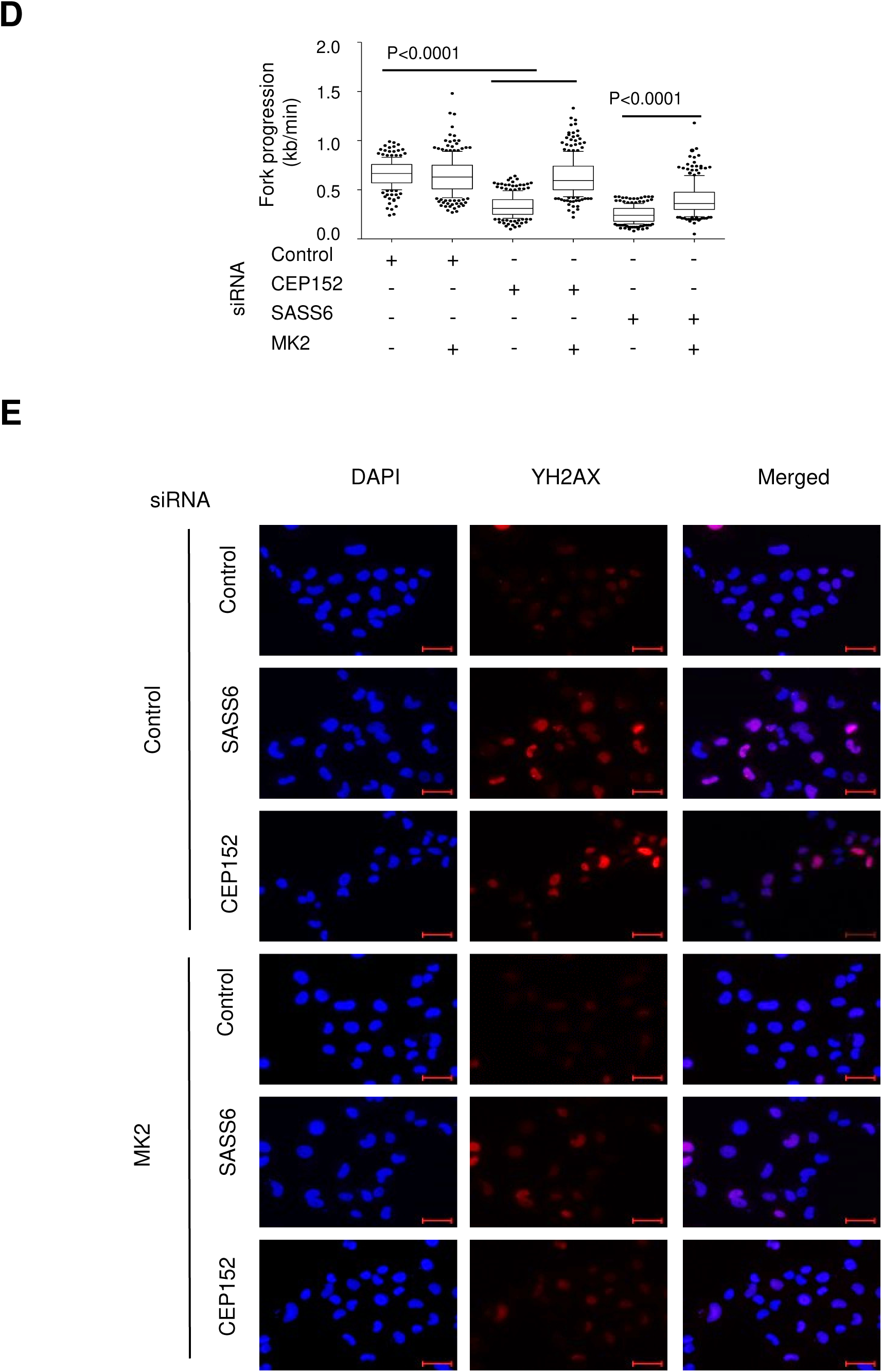
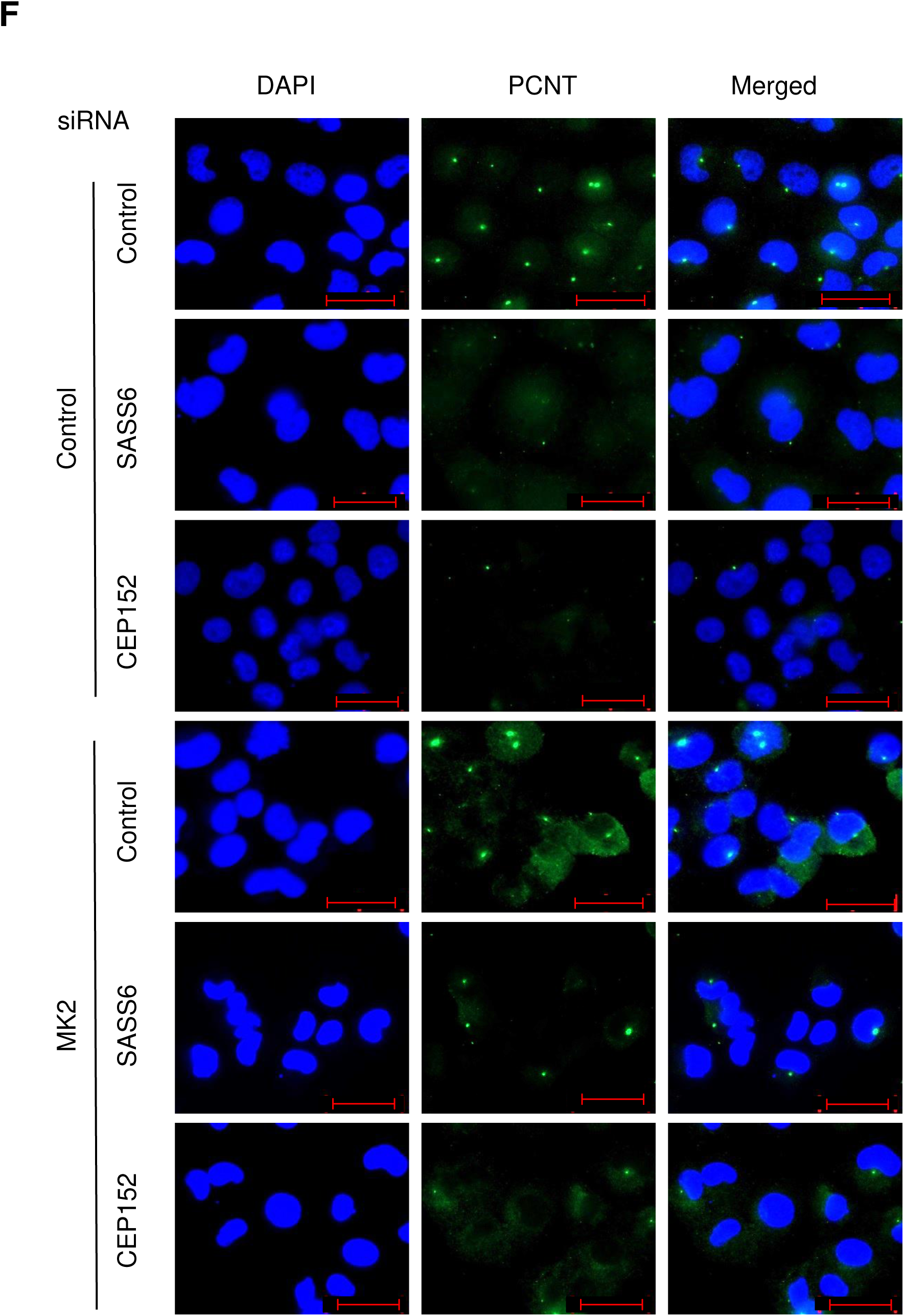
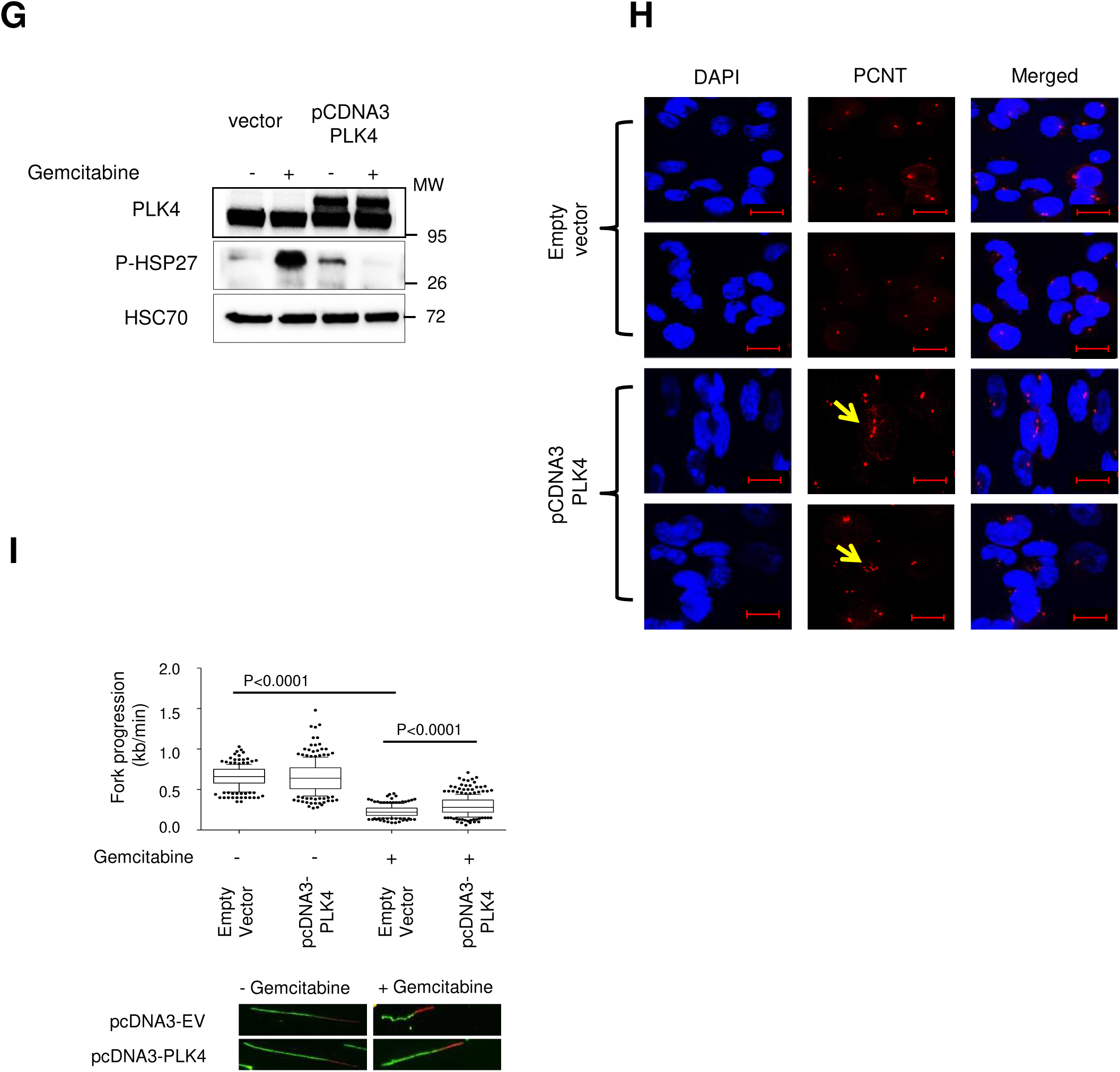
**(A)** MK2 inhibition rescues replication fork progression upon treatment with Centrinone. Synchronized H1299 cells were treated as in Figure 4 B. **(B)** MK2 inhibition diminishes the accumulation of γH2AX upon PLK4 inhibition. Images corresponding to Figure 4 C. **(C)** MK2 inhibition does not detectably affect the number of centrosomes upon PLK4 inhibition, as detected by immunostaining of PCNT. Images corresponding to Figure 4 E. **(D)** Knockdown of MK2 rescues DNA replication in cells that were depleted of centrosomal components. Synchronized H1299 cells were treated as in Figure 4 G. **(E)** Knockdown of MK2 largely abolishes the accumulation of γH2AX upon depletion of centrosomal components. Images corresponding to Figure 4 H. **(F)** Knockdown of MK2 does not detectably affect the numbers of centrosomes when centrosomal components are depleted. Images corresponding to Figure 4 J. **(G)** PLK4 overexpression diminishes MK2 activation in the presence of gemcitabine. Immunoblot analysis to confirm PLK4 overexpression (note that the apparent molecular weight is increased due to the Flag tag). MK2 activity, as revealed by HSP27 phosphorylation, is increased by gemcitabine, but not when PLK4 is overexpressed. **(H)** Increased centrosome formation upon PLK4 overexpression. Synchronized H1299 cells were subjected to plasmid transfection (empty pcDNA3 or pcDNA3-PLK4) for 48 hrs. Centrosomes were detected by immunostaining PCNT, and the DAPI signal was used to detect the nuclei. Scale bar represents 20 μm. Arrows, amplified centrosomes in response to overexpressed PLK4. **(I)** PLK4 overexpression partially rescues DNA replication in gemcitabine-treated cells. Synchronized H1299 cells were subjected to plasmid transfection (pcDNA3, pcDNA3-PLK4) for 48 hrs, followed by treatment with 300nM Gemcitabine for 2 hrs prior to harvesting. DNA replication fork progression was determined in fiber assays.

**Supplementary Figure 5, corresponding to Figure 5.**
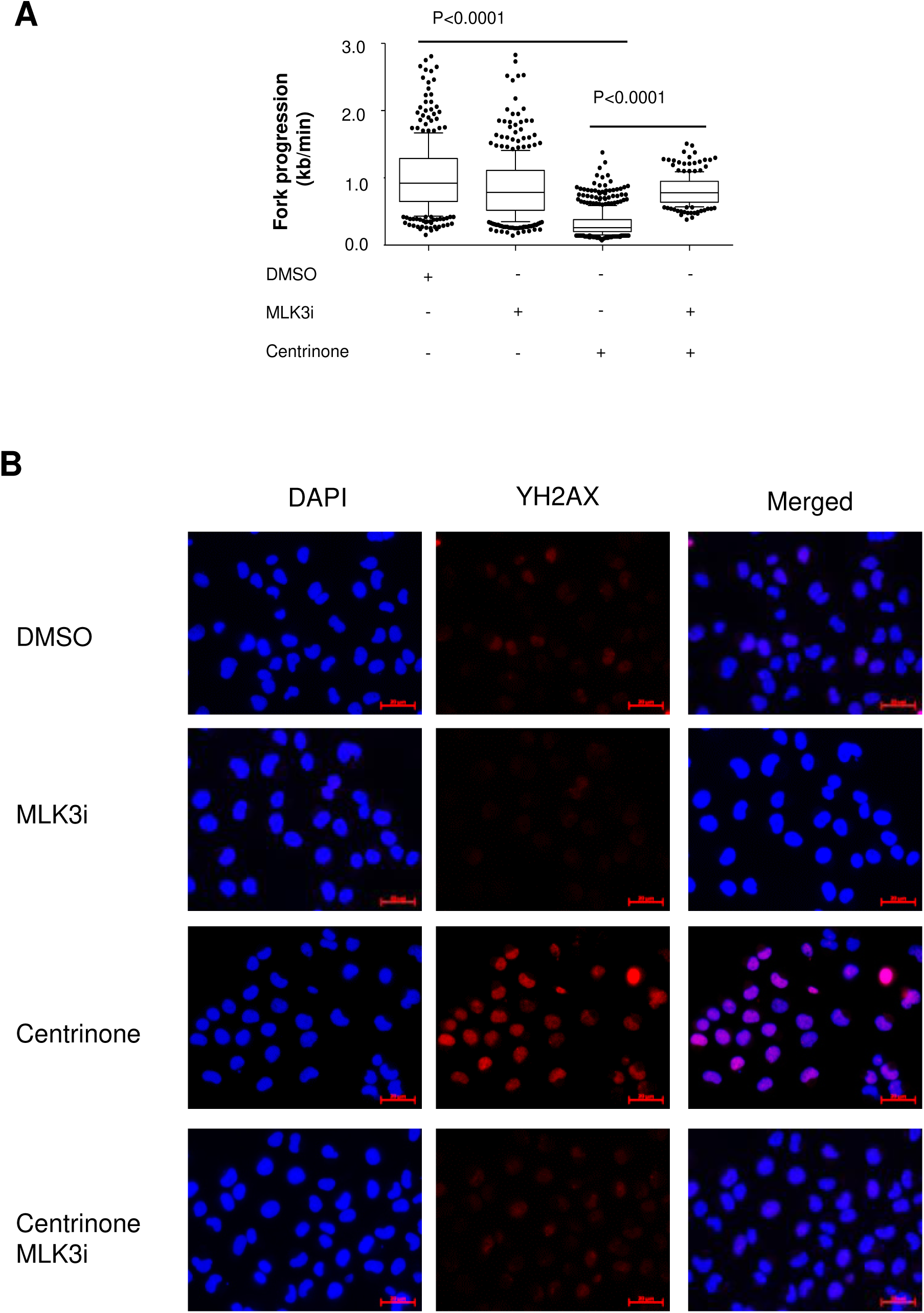
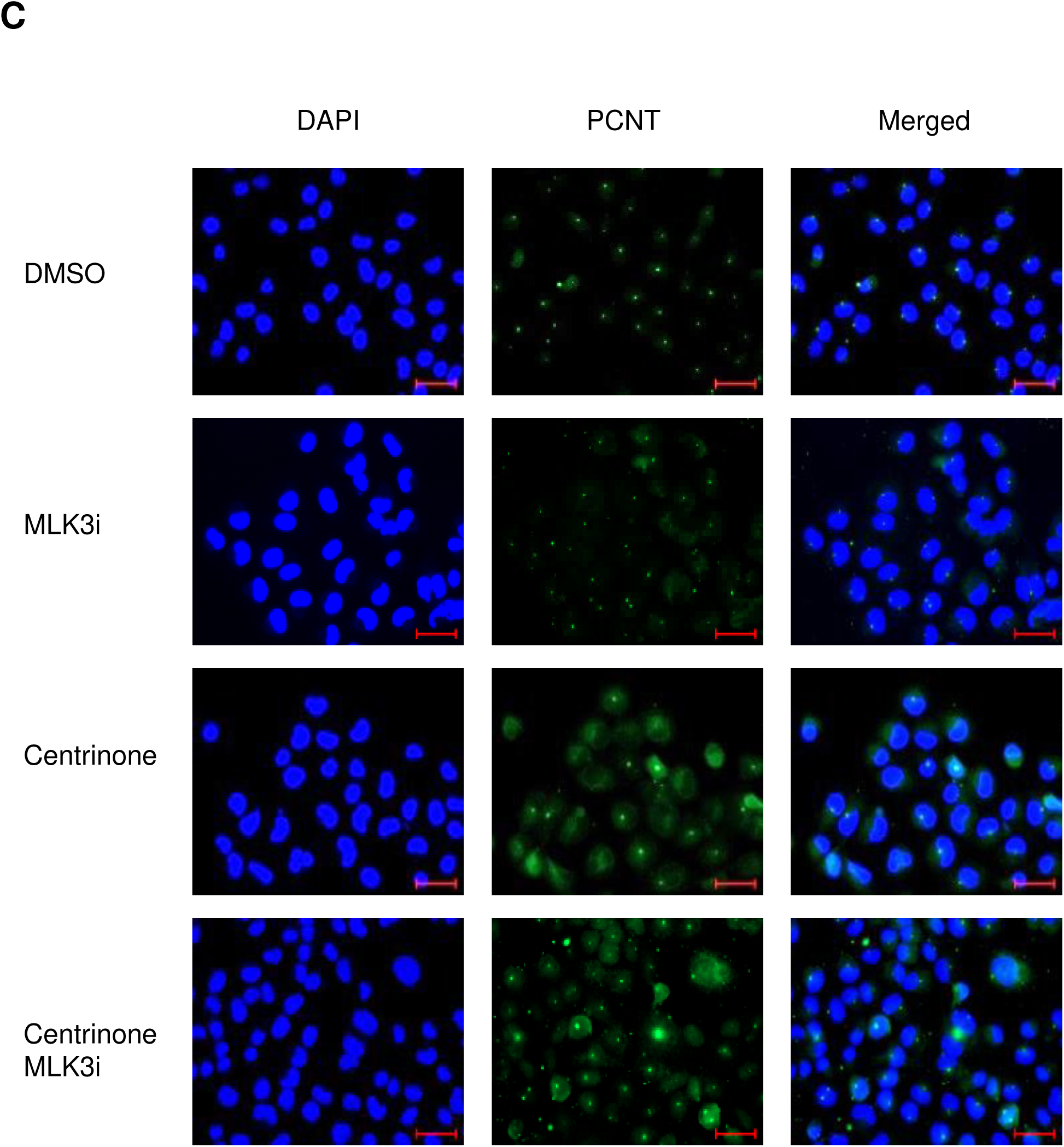
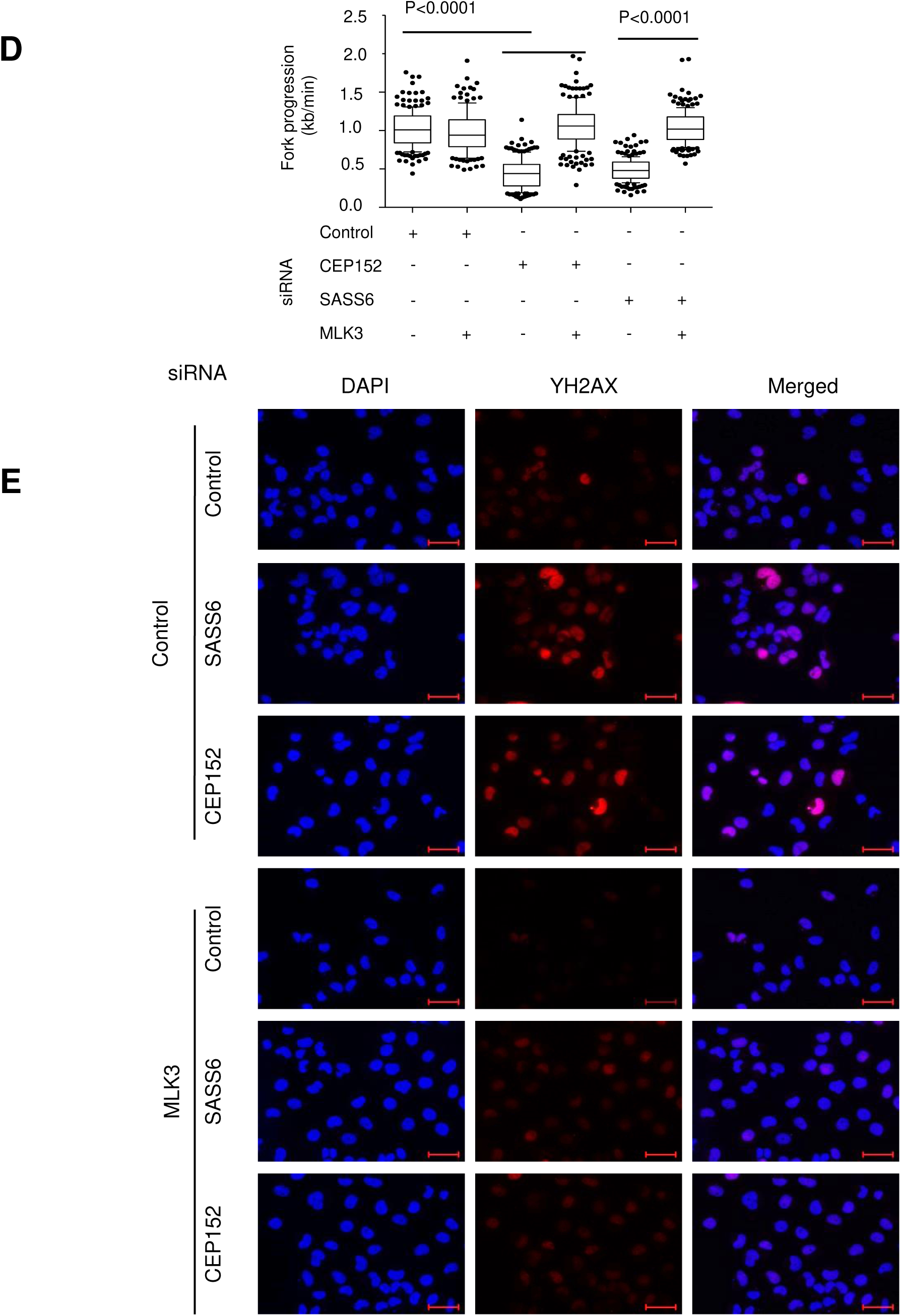
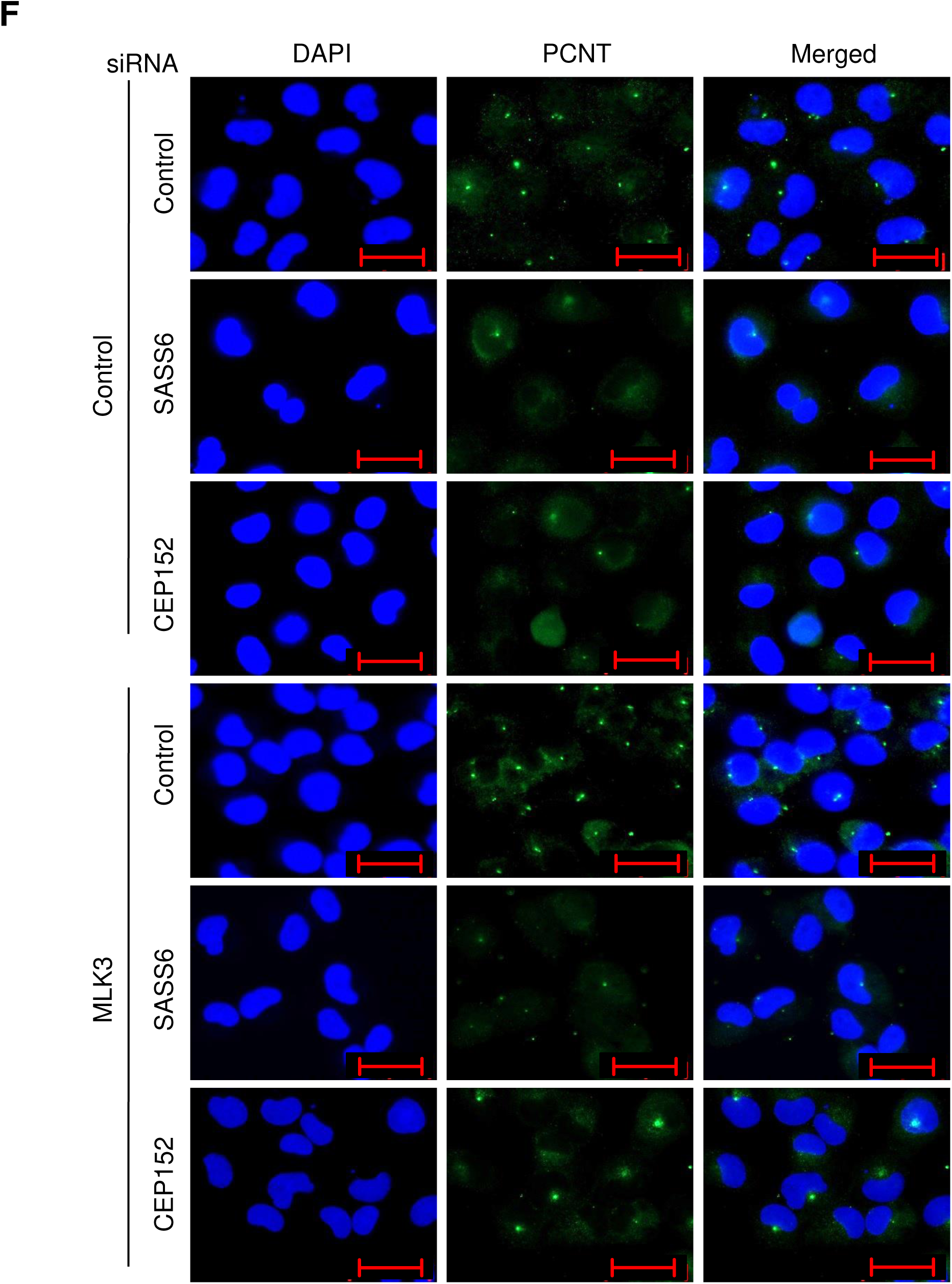
**(A)** MLK3 inhibition rescues DNA synthesis upon PLK4 inhibition. Synchronized H1299 cells were treated as Figure 5 D. **(B)** MLK3 inhibition diminishes the accumulation of γH2AX in the context of PLK4 inhibition by Centrinone. Images corresponding to Figure 5 E. **(C)** MLK3 inhibition partially restores the number of centrosomes upon PLK4 inhibition. Images corresponding to Figure 5 F. **(D)** MLK3 depletion restores DNA replication upon knockdown of centrosomal components. Synchronized H1299 cells were treated for fiber assays as in Figure 5 H. **(E)** Knockdown of MLK3 reduces the accumulation of γH2AX when centrosomal components are depleted. Images corresponding to the quantification in Figure 5 I. **(F)** Depleting MLK3 partially restores detectable centrosomes upon knockdown of CEP152 (but not SASS6). Images corresponding to the quantification shown in Figure 5 J.

**Supplementary Figure 6, corresponding to Figure 6.**
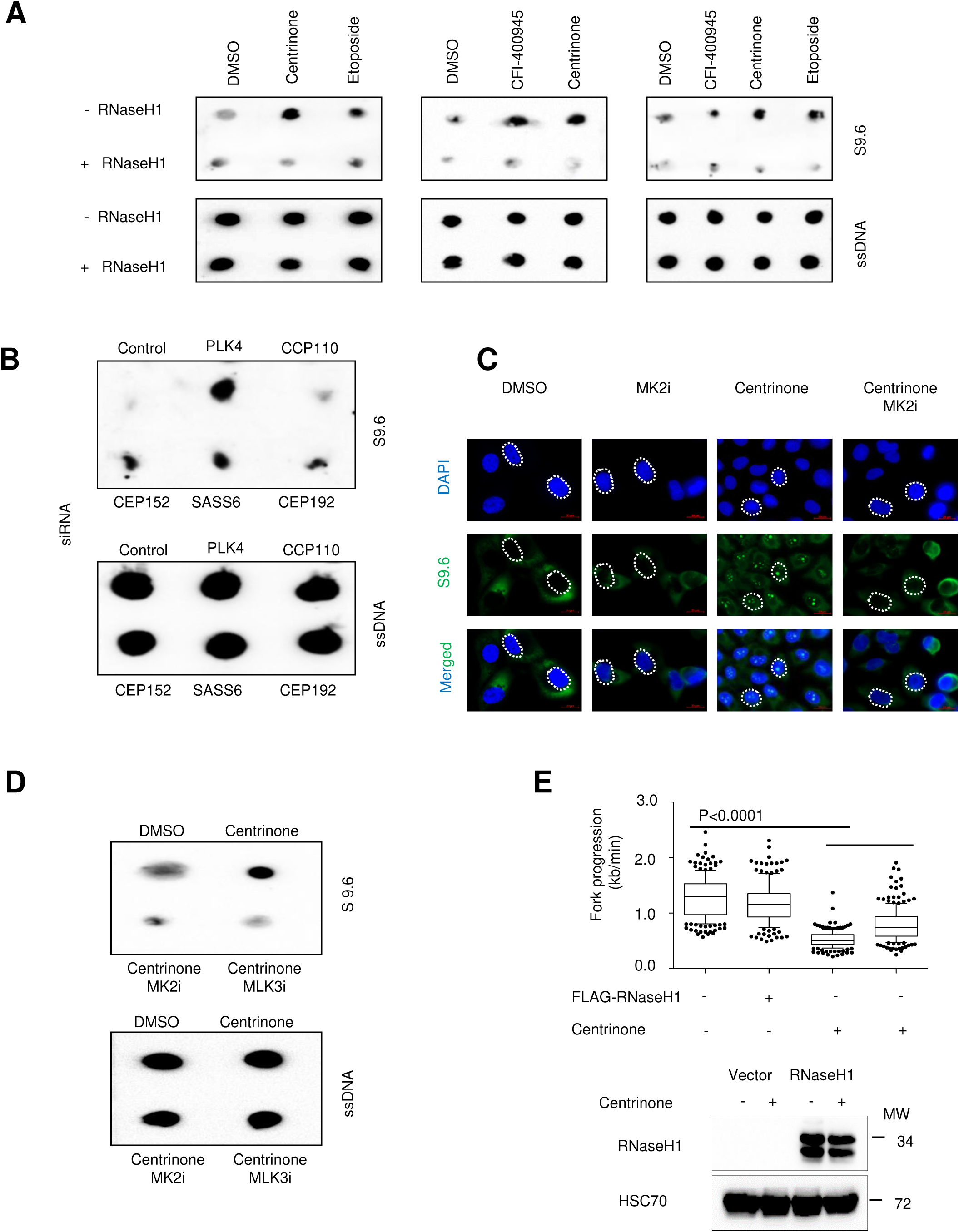
**(A)** Dot blots showing the accumulation of RNA:DNA hybrids upon PLK4 inhibition by Centrinone or CFI-400945. The experiment was carried out as in Figure 6 C. **(B)** Dot blot showing the accumulation of RNA:DNA hybrids upon depletion of centrosome components, corresponding to the quantification shown in Figure 6 E. **(C)** Immunofluorescence images showing the formation of R loops formation upon PLK4 inhibition, along with their rescue by MK2 inhibition, corresponding to the quantification in Figure 6 F. **(D)** Dot blot reflecting the accumulation of RNA:DNA hybrids upon PLK4 inhibition, and their reduction by inhibition of MK2 or MLK3, corresponding to the quantification shown in Figure 6 G. **(E)** RNaseH1 rescues DNA replication in the context of PLK4 inhibition. Biological replicates for the fiber assays presented in Figure 6 H.

**Supplementary Figure 7, corresponding to Figure 7.**
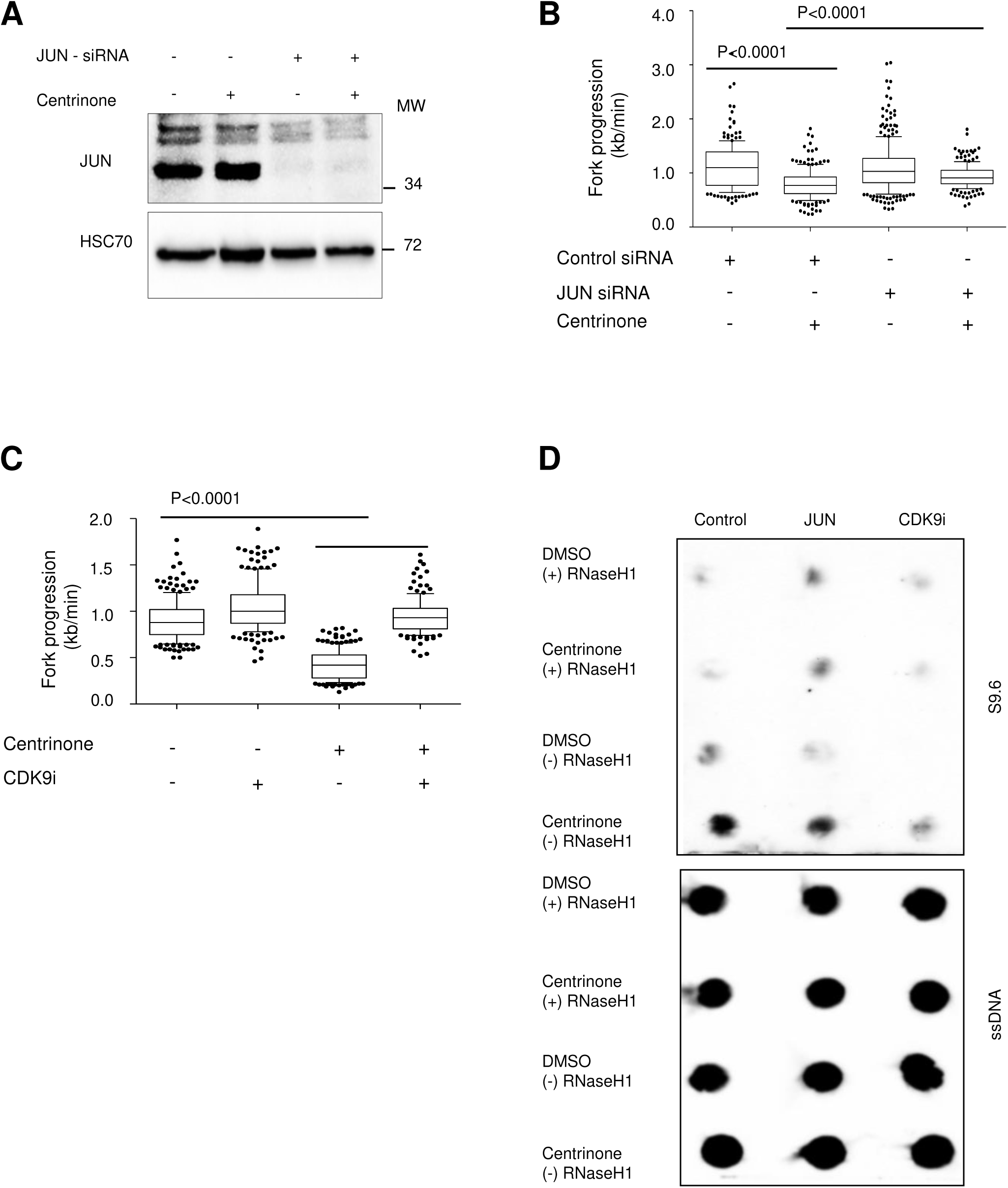
**(A)** Validation for JUN knockdown in H1299 cells by immunoblot analysis. HSP70 staining was performed as a loading control. In the control experiments, scrambled siRNA was transfected instead of siRNA targeting JUN. **(B)** JUN depletion partially rescues DNA replication upon cell treatment with Centrinone. Synchronized H1299 cells treated as Figure 7 D **(C)** CDK9 inhibition rescues DNA replication upon treatment with a PLK4 inhibitor. Synchronized H1299 cells treated as Figure 7 E **(D)** JUN depletion reduces RNA:DNA hybrids in the context of PLK4 inhibition, whereas CDK9 inhibition abolishes them. Dot bot membrane corresponding to the quantification in Figure 7 F

**Supplementary Figure 8, corresponding to Figure 8.**
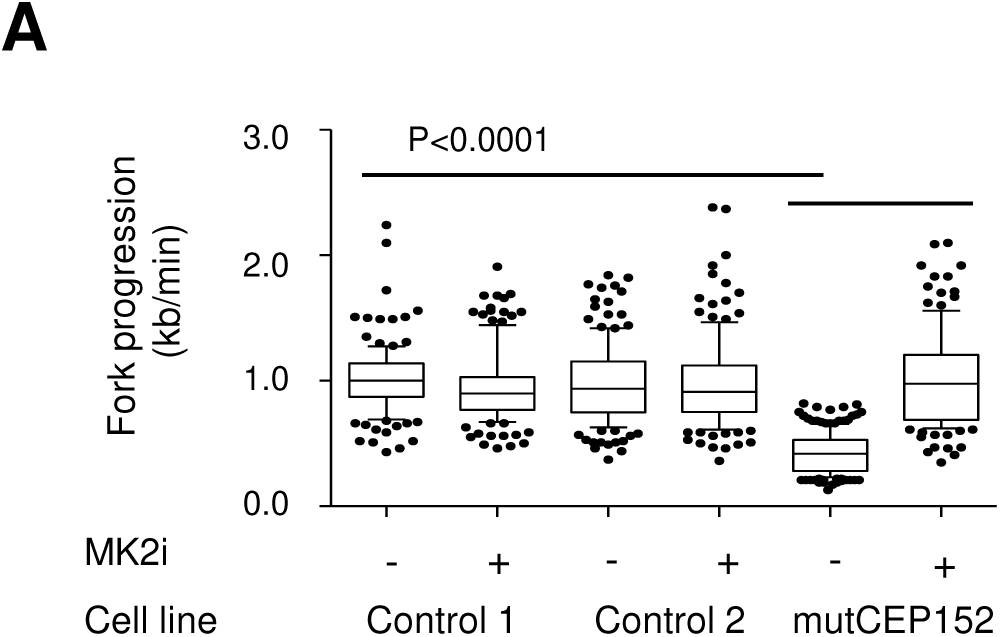
**(A)** MK2 inhibition restores DNA replication in Fibroblasts with mutant CEP152. Replicate to Figure 8 A.

**Supplementary Table 1**

This table is listing the materials used in all experiments, including antibodies, siRNAs, and plasmids, with their sources and identifier codes.

**Supplementary Table 2**

Raw data obtained by measuring the DNA replication fork progression using fiber assays. Data corresponding to results presented in Figures 1-8. The length of the fibers was measured using ImageJ software, while the box blot graphs and statistical analyses were made using the Prism software. ∼ 200 fibers were measured per treatment scheme for each experiment.

**Supplementary Table 3**

The raw data of cell confluence measured using the Celigo Cytometer. Data corresponding to results presented in Figure 4 D and 4 I and Figure 8 D. Cells were treated as stated in the legend to each of these figures. Cell confluence was measured every 24 hrs for up to 7 days. Experiments were carried out in 2 biological replicates with 3 technical replicates each, for every time point.

